# Parallel evolution of local adaptation in *Littorina saxatilis* inferred with whole-genome pool-seq data

**DOI:** 10.1101/2023.09.18.558243

**Authors:** João Carvalho, Rui Faria, Roger K. Butlin, Vítor C. Sousa

**Author notes:** Corresponding author’s.

## Abstract

Parallel evolution of phenotypic divergence offers compelling evidence supporting the influence of natural selection on the divergence process. However, the existence of ecotypes adapted to different habitats in separate locations is not a definitive proof of parallel evolution. Here, we leverage a large pool-seq dataset of the rocky-shore gastropod, *Littorina saxatilis* to compare and contrast two explicit scenarios of ecotype formation. Populations of this species inhabit contrasting habitats, where the main selective pressure is either wave action or crab predation. Using approximate Bayesian computation to jointly infer demographic parameters and account for pool-seq specific sources of uncertainty, our analysis reveals that ecotype formation at a large geographical scale has occurred in parallel. Parameter estimates provide strong support for a demographic history marked by the spatial separation of the ancestral populations that later gave rise to parallel evolution of ecotypes in Spain and Sweden. Additionally, ecotype formation occurred in the face of continuous gene flow between the diverging ecotypes. These results shed new light on a important model of speciation driven by ecological factors and emphasize the effectiveness of combining pool-seq with ABC for studying parallel evolution across diverse geographical regions.

## INTRODUCTION

Understanding the processes involved in the adaptation of populations to different environments is one of the key goals in evolutionary biology (Seehausen et al., 2014). Divergent natural selection can drive local adaption of populations to different ecological conditions. Importantly, the action of divergent natural selection in distinct environments is a common driving force behind population divergence and the evolution of reproductive isolation, which can ultimately result in new species (Nosil, 2012; Schluter, 2009). By uncovering the evolutionary drivers of adaptive divergence, we can improve our understanding of how biodiversity is generated and maintained, both within and between species. Gene flow between locally adapted populations may oppose local adaptation and the ensuing divergence process (Akerman & Bürger, 2014; Lenormand, 2002). Thus, the spatial context of population divergence and the extent (or possibility) of gene flow play a crucial role in determining the outcome of the speciation process (Smadja & Butlin, 2011). Therefore, a complete understanding of local adaptation and the evolution of reproductive isolation can only be obtained by considering the biogeographical and demographic history of populations (He et al., 2019; Hewitt, 2011). Genomic data and coalescent-modeling approaches have been used to infer such historical sequences of events (Sousa & Hey, 2013). For instance, a recent study by Portinha et al. (2022) used genomic data from wood ant species and coalescent-based modeling to reconstruct their demographic history, uncovering a pattern of divergence with continuous asymmetrical gene flow. Similarly, demographic analysis using whole-genome sequence data of two sympatric stickleback species showed that a long period of allopatry might have promoted the evolution of intrinsic incompatibilities (Yamasaki et al., 2020).

The action of divergent natural selection occurring in multiple populations facing similar environmental conditions can lead to similar phenotypic traits, a phenomenon known as parallel evolution (Bolnick, Barrett, Oke, Rennison, & Stuart, 2018; Elmer & Meyer, 2011). This phenotypic parallelism can be explained by different processes, ranging from convergent evolution acting on *de novo* mutations arising in different populations, to selection based on standing genetic variation leading to both phenotypic and genomic parallelism. Given the opposing effect of gene flow and the randomness of other processes such as genetic drift, parallel evolution stands as one of the most convincing forms of evidence for the significant and predictable role of natural selection in driving adaptive evolution (Faria et al., 2014; Nosil, 2012). Moreover, when adaptation to similarly divergent pairs of environments leads to the repeated evolution of reproductive isolation (i.e., parallel speciation), it underscores the potential of natural selection in driving the speciation process (Johannesson, 2001). However, environments can differ along multiple axes, potentially leading replicate populations to evolve along trajectories that deviate from the parallel expectation and vary not only in the magnitude but even in the direction of phenotypic divergence (Langerhans, 2018; Oke, Rolshausen, LeBlond, & Hendry, 2017). Additionally, parallelism at the phenotypic level does not equate to parallelism at the genomic level given that different genetic changes can underlie the same phenotypic solution (Bainbridge et al., 2020; Poore et al., 2022).

Examples of parallelism at the phenotypic level associated with varying degrees of parallelism at the genomic level have been found in populations of the beach mouse, *Peromyscus polionotus* (Steiner, Römpler, Boettger, Schöneberg, & Hoekstra, 2009) and the American lake whitefish, *Coregonus clupeaformis* (Laporte et al., 2015). Another noteworthy example is the marine three-spine stickleback, *Gasterosteus aculeatus*, that has independently colonized many freshwater sites. At each of those sites, the same pattern of reduced armour evolution has been observed (Colosimo et al., 2005; Jones et al., 2012). This pattern is accompanied by parallelism at the underlying major quantitative trait locus (EDA; Colosimo et al. 2005), representing a textbook example of phenotypic and genomic parallelism. Despite this, there are exceptions to this parallelism (Fang, Kemppainen, Momigliano, Feng, & Merilä, 2020), with different levels of lateral plating observed (Hansson, Fischer, Mazzarella, Voje, & Vøllestad, 2016) and the absence of genomic parallelism in the reduction of pelvic armour in replicate freshwater populations (Chan et al., 2010). Such deviations can be explained by non-adaptive processes such as genetic drift or differences in gene flow between habitats (Ferchaud & Hansen, 2016; Hendry & Taylor, 2004) or they might reflect adaptive responses to habitat-specific differences among similar habitats (DeFaveri, Shikano, Shimada, Goto, & Merilä, 2011). Interestingly, results from a meta-analysis suggest that the probability of genomic parallelism depends on the age of the populations and their access to the same pool of standing genetic variation (Conte, Arnegard, Peichel, & Schluter, 2012). Indeed, genomic parallelism is expected to be more prevalent in populations that share a common ancestry and are in close geographic proximity (Bohutínská et al., 2021; Rennison, Delmore, Samuk, Owens, & Miller, 2020). This is not surprising because populations with a recent shared evolutionary history and higher connectivity have access to a common pool of standing genetic variation. Conversely, habitat specific changes in the direction of selection or in the selective pressure itself can lead to nonparallel patterns of phenotypic divergence (Kaeuffer, Peichel, Bolnick, & Hendry, 2012; Thurman et al., 2023).

Interpreting the pattern of parallel local adaptation in the presence of contemporary gene flow is not straightforward since it can arise as a consequence of markedly different historical sequences of events (Johannesson et al., 2010). In particular, two broadly contrasting scenarios can give rise to this parallel pattern. In one scenario, the initial adaptive divergence evolves once (in sympatry or allopatry), with subsequent colonization of similar pairs of environments by differentially adapted forms. Alternatively, divergence occurs repeatedly and independently in multiple localities (Butlin et al., 2014; Johannesson et al., 2010). Interestingly, even if these repeated origins of phenotypic divergence are independent, there is a possibility that the same adaptive alleles are used, possibly originating from standing genetic variation or gene flow across localities. The first step towards distinguishing between these alternatives requires the use of putatively neutral genetic markers to establish the demographic history of the populations. This can be achieved by using genomic data to inform coalescent-based models of parallel local adaptation integrated in an model-based inference framework such as Approximate Bayesian Computation (ABC). However, this would require sampling and sequencing multiple populations across a wide geographical range, which might quickly become cost prohibitive. A cost-effective approach is to pool DNA from multiple individuals and sequence them together (pool-seq), which makes it possible to quantify and characterize the genetic variation within and between populations (Schlötterer, Tobler, Kofler, & Nolte, 2014). However, estimates of allele frequency obtained with pool-seq can be affected by variations in DNA concentration of the pooled individuals and amplification/sequencing efficiency, on top of more common sources of errors such as depth of coverage variation across sites and sequencing errors. Nevertheless, pool-seq allows for the analysis of more individuals with similar or more precise allele frequency estimates compared to individual-based sequencing, making it suitable for inferring population genetic structure and estimating effective population sizes and divergence times (Collin et al., 2021; Futschik & Schlötterer, 2010).

Despite a general awareness that different sequences of events can lead to the same pattern of parallel divergence, relatively few studies have explicitly tested the parallel origin scenario against alternative hypotheses. Here we apply a recently developed approach (Carvalho, Morales, Faria, Butlin, & Sousa, 2023) to a large pool-sequencing dataset of the rough periwinkle, *Littorina saxatilis*, to contrast alternative scenarios for the origin of locally adapted ecotypes in this species. This rocky-shore gastropod has a wide distribution, covering most of the North Atlantic despite having a low lifetime dispersal (Reid, 1996). Across the species distribution, two ecotypes can be found in close proximity: a smaller morph with a thin shell and a larger aperture that allows it to withstand heavier wave exposure and a larger morph with a thicker shell and a small aperture that confers more protection against crab predation (Johannesson et al., 2010). These ecotypes have been the focus of numerous studies (e.g. Conde-Padín, Cruz, Hollander, and Rolan-Alvarez 2008; Grahame, Wilding, and Butlin 2006) and evidence suggests that sexual selection and assortative mating are also involved in reproductive isolation (Perini, Rafajlović, Westram, Johannesson, & Butlin, 2020).

Recently, multiple chromosomal rearrangements associated with the ecotypes (Faria et al., 2019) have been connected to variation linked with adaptive traits (Koch et al., 2021). However, some studies indicate that loci outside inversions may also contribute to the divergence (Kess & Boulding, 2019; Koch, Ravinet, Westram, Johannesson, & Butlin, 2022; Westram, Faria, Johannesson, & Butlin, 2021). A common thread of most *L. saxatilis* studies is their focus on only one of three European regions: Galicia in northwest Spain, the west coast of Sweden and the northeast coast of England. An independent and parallel origin of these ecotypes has often been invoked (Quesada, Posada, Caballero, Morán, & Rolán-Alvarez, 2007). However, some of these arguments were questioned (Butlin, Galindo, & Grahame, 2008) and it has been found that genomic parallelism is limited (Koch et al., 2022; Ravinet et al., 2016). Furthermore, the Iberian populations share a demographic history that is clearly different from the remaining European populations (Blakeslee et al., 2021; Morales et al., 2019; Panova et al., 2011), while the Swedish and British populations have probably shared a post-glacial colonization history. A single study (Butlin et al., 2014) has examined and compared different models of the demographic history of the ecotypes across a wide geographical range. However, this study relied on a limited set of genetic data, specifically mitochondrial and nuclear DNA sequences combined with amplified fragment length polymorphism data. Building upon previous research, we use whole-genome data to compare and contrast two explicit scenarios of ecotype formation. Our aim is to determine whether *L. saxatilis* ecotype formation on a large scale can be attributed to a single event or a series of parallel events.

## MATERIAL AND METHODS

### Models of ecotype formation: single vs. parallel

We compared two contrasting scenarios of ecotype formation (Figure 1). We represented these scenarios using a four-population model, wherein the four extant populations correspond to the two distinct ecotypes observed at two separate geographical locations. We considered 32 parameters of interest for both scenarios. These include the size of the four present-day populations, denoted as *N*_1_ to *N*_4_, as well as the two ancestral populations, *NA*_1_ and *NA*_2_. Additionally, the time of the recent split events between populations in the first location (*RS*_1_) and between populations in the second location (*RS*_2_), as well as the ancient split event (*T_As_*) are all measured in generations. We also considered migration rates between the divergent ecotypes at each location, with *m*_12_ and *m*_21_ representing the migration from *N*_1_ to *N*_2_ and from *N*_2_ to *N*_1_, respectively. Similarly, the migration rates *m*_34_ and *m*_43_ represent the migration (forward in time) from *N*_3_ to *N*_4_, and from *N*_4_ and *N*_3_, respectively. We also considered migration between similar ecotypes inhabiting different locations and between ancestral populations. Note that to avoid increasing the number of parameters under consideration, the migration rates between similar ecotypes and between ancestral populations were assumed to be identical regardless of the direction. However, the scaled migration rate 4*Nm* can be different. In order to estimate the timing of events, we considered as a parameter the time interval between the older recent split event and the ancient split (Δ*_s_* = *T_As_ − RS*). We incorporated the existence of bi-directional or uni-directional barrier loci in our model. For the bi-directional barrier, we included a parameter (*P_no_*) to control the proportion of loci where there was no migration in either direction between the divergent ecotypes. As for the uni-directional barrier, we included parameters *P_cw_*and *P_wc_* representing the proportion of loci where no migration occurred from Crab to Wave or from Wave to Crab, respectively. We assumed that these parameters (*P_no_*, *P_cw_*, and *P_wc_*) were the same across both localities. Depending on the topology, the four-population model can represent either a single origin scenario where ecotypes originate in distinct locations before dispersing and colonizing two geographic areas (Figure 1A), or a parallel origin scenario where each location is colonized independently, leading to the independent divergence of the distinct ecotypes (Figure 1B).

**Figure 1:**
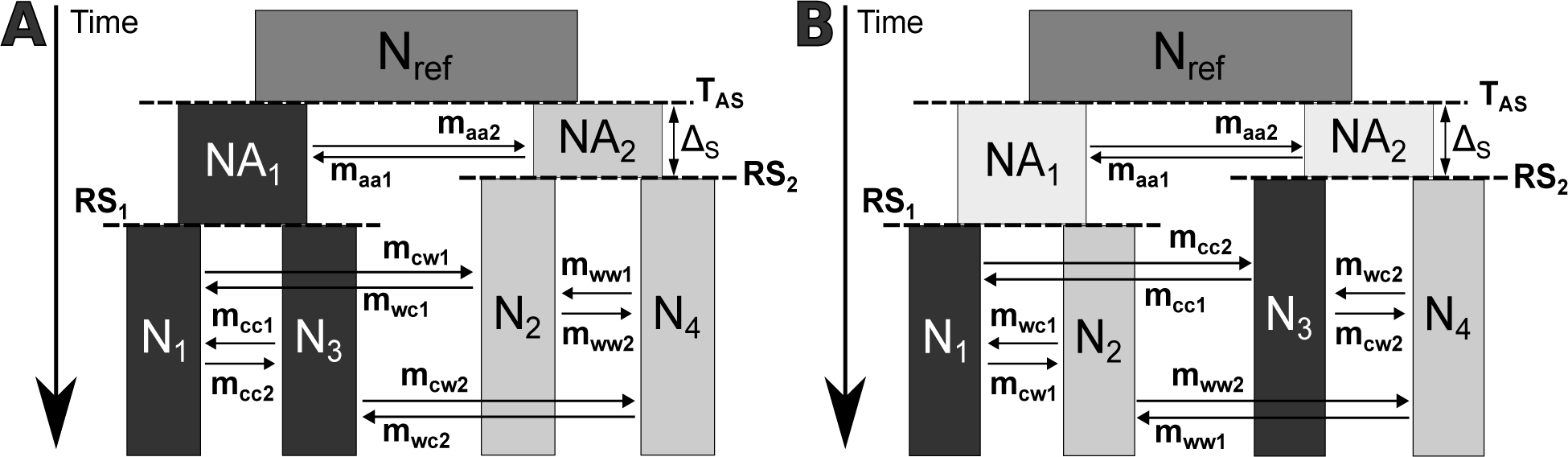
Demographic models for the single (A) and parallel (B) ecotype formation scenarios. Dark shading indicates one of the ecotypes, light shading the other ecotype. Parameters used were: *N_re f_* - effective size of the ancestral population, *NA*_1_ and *NA*_2_ - size of the two ancestral populations, *N*_1_ - *N*_4_ - sizes of the present-day populations, *RS*_1_ - time of separation of the present-day populations at the first location (in generations), *RS*_2_ - time of separation of the present-day populations at the second location (in generations), *T_As_* - time of the ancient split event (in generations), Δ*_s_* - time interval between the two split events (in generations), *m_cw_*_1_ - probability per generation that an individual migrates from Crab *N*_1_ to Wave *N*_2_ (forward in time), *m_wc_*_1_ - probability per generation that an individual migrates from Wave *N*_2_ to Crab *N*_1_ (forward in time), *m_cw_*_2_ - probability per generation that an individual migrates from Crab *N*_3_ to Wave *N*_4_ (forward in time), *m_wc_*_2_ - probability per generation that an individual migrates from Wave *N*_4_ to Crab *N*_3_ (forward in time), *m_cc_*_1_ - probability per generation that an individual migrates from Crab *N*_3_ to Crab *N*_1_ (forward in time), *m_cc_*_2_ - probability per generation that an individual migrates from Crab *N*_1_ to Crab *N*_3_ (forward in time), *m_ww_*_1_ - probability per generation that an individual migrates from Wave *N*_4_ to Wave *N*_2_ (forward in time), *m_ww_*_2_ - probability per generation that an individual migrates from Wave *N*_2_ to Wave *N*_4_ (forward in time), *m_aa_*_1_ - probability per generation that an individual migrates from *NA*_2_ to *NA*_1_ (forward in time) and *maa*2 - probability per generation that an individual migrates from *NA*_1_ to *NA*_2_ (forward in time).

### Simulation of pool-seq data

We simulated gene trees using coalescent theory and the *scrm* simulator (Staab, Zhu, Metzler, & Lunter, 2015). Mutations followed the infinite sites model, with a mutation rate (*µ*) per site per generation. Gene trees were simulated for each locus with the same sample size as the number of individuals in the pool, which corresponds to 100 diploid individuals (200 haplotypes) for each population. During gene tree simulations, the actual haplotypes of all individuals in the pool were assumed to be known, and the effect of pooling was simulated later. To simulate genotypes, random mating was assumed within each population, pairing haplotypes randomly at each locus to obtain genotypes for each biallelic SNP.

To model allele frequencies at biallelic SNPs obtained with pool-seq we followed a series of steps detailed in Carvalho et al. (2023). We simulated pool-seq data using parameters obtained from real *L. saxatilis* pool-seq data (see below). Briefly, we modelled the depth of coverage at each SNP (number of reads per site) using a negative binomial distribution (Sampson, Jacobs, Yeager, Chanock, & Chatterjee, 2011). This distribution is defined by the mean and variance of the coverage of the real data. DNA extraction for the *L. saxatilis* pool-seq data was performed for batches of five individuals by combining foot muscle tissue from five snails in one tube (Morales et al., 2019). The DNA from each batch was then pooled to create the final pools for sequencing. Thus, for each SNP, we modelled the possibility of uneven contributions between those batches of five individuals. This uneven contribution was modeled as a multinomial-Dirichlet distribution, where the number of reads from each batch is given by a multinomial with expected proportion of reads from each batch following a Dirichlet distribution.

We then modelled the heterogeneity in the contribution of each individual within each batch, following the same approach: number of reads from each individual as a multinomial distribution, where the expected proportion of reads from each individual followed a Dirichlet distribution. In both cases, the dispersion of the Dirichlet distribution is determined by explicit pool-seq error parameters (Carvalho et al., 2023; Gautier et al., 2013). These parameters influence the variance of the proportion of reads contributed by each batch or individual, representing the random variation in the batch or individual contribution to the pool (Carvalho et al., 2023; Gautier et al., 2013). Higher values of the pool-seq error parameter lead to increased variance, resulting in greater heterogeneity in the contributions of both individual and batches. This model assumes an equal expected contribution of reads from all individuals, with errors arising from unequal contributions accounted for through dispersion parameters that affect the variance. To model the potential impact of sequencing and mapping errors, we considered the possibility that the allele observed in the pool’s reads might not corresponded to the true genotype of the sampled individuals. For each individual, the count of reads with the alternative allele was modeled using a binomial distribution. This distribution assumed that, with a given error rate, the reference allele could be incorrectly identified as the alternative allele, or vice versa. We then defined which of the two alleles (i.e., reference or alternative) corresponded to the minor-allele and applied a minor-allele filter, discarding all SNPs with less than two minor-allele reads. We did not consider sequencing errors occurring at invariant sites, as the simulated pool-seq data only included polymorphic sites. Furthermore, any such errors would probably be eliminated by the minor-allele filter. To mimic real *L. saxatilis* pool-seq data (see below), we applied a coverage-based filter, removing all sites with a coverage below 14x or above 204x. Finally, we estimated the minor-allele frequencies as the proportion of reads with the minor-allele.

### ABC implementation

Our ABC implementation, using a rejection algorithm, followed several steps (Carvalho et al., 2023). In short, we started by sampling demographic and pool-seq parameters from prior distributions (Table 1). Subsequently, we simulated genotypes for each individual at *L* loci using coalescent theory and the sampled demographic history parameters. We then simulated pool-seq data for each biallelic SNP and implemented filters based on depth of coverage and minor-allele counts. Afterwards, we computed summary statistics for both the observed and simulated data and calculated the Euclidean distance between the observed and simulated summary statistics, ensuring standardization so that that all summary statistics have the same mean and variance. Next, we rejected parameters that led to summary statistics with distances exceeding a specified tolerance threshold (i.e., rejection step). The accepted parameters from the rejection step are used to approximate the posterior distribution of the parameters. Finally, we applied a post-processing regression to adjust the accepted parameter values (Beaumont, Zhang, & Balding, 2002).

**Table 1:**
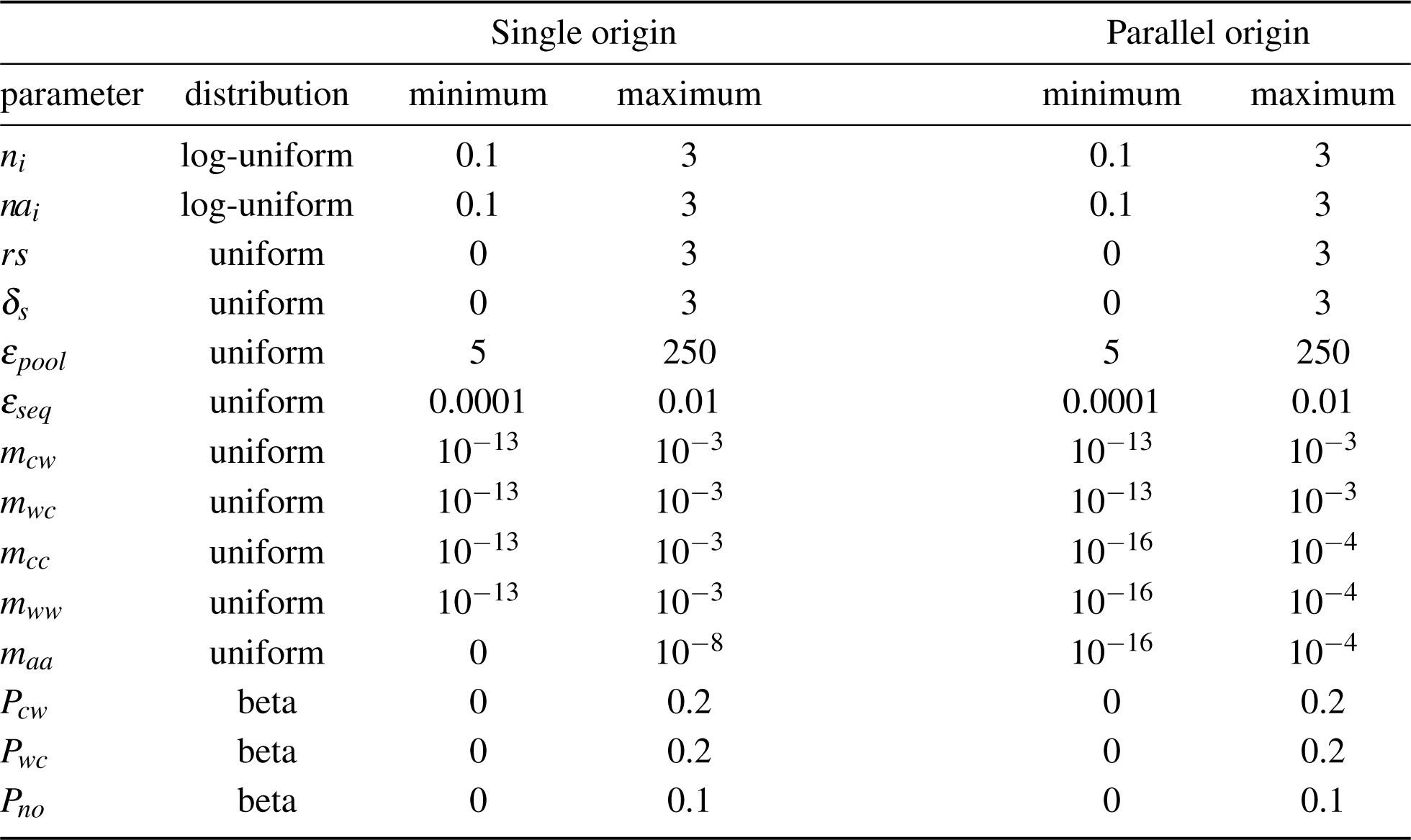
Prior distributions and their ranges for each parameter. Parameters are presented for the single and parallel origin scenarios. *n_i_* - relative sizes of the extant populations (*n*_1_*, n*_2_*, n*_3_*, n*_4_); *na_i_* - relative sizes of the ancestral populations (*na*_1_*, na*_2_); *rs* - relative time of the recent split event; *δ_s_* - relative time interval between *rs* and the ancient split event (*t_As_*); *ε_pool_* - experimental error introduced by the pooling procedures; *ε_seq_*- error associated with sequencing and mapping errors; *m_cw_* - probability per generation that an individual migrates from the the Crab population to the Wave population (forward in time), *m_wc_*- probability per generation that an individual migrates from the Wave population to the Crab population (forward in time); *m_cc_* - probability per generation that an individual migrates between Crab populations from different geographic locations (forward in time); *m_ww_* - probability per generation that an individual migrates between Wave populations from different geographic locations (forward in time); *m_aa_* - probability per generation that an individual migrates between ancestral populations (forward in time); *P_cw_* - proportion of the simulated loci where no migration occurs from the Crab to the Wave population; *P_wc_* - proportion of the simulated loci where no migration occurs from the Wave to the Crab population and *P_no_*- proportion of the simulated loci where no migration occurs between ecotypes.

| parameter | distribution | Single origin |  | Parallel origin |  |
| --- | --- | --- | --- | --- | --- |
|  |  | minimum | maximum | minimum | maximum |
| $n_i$ | log-uniform | 0.1 | 3 | 0.1 | 3 |
| $na_i$ | log-uniform | 0.1 | 3 | 0.1 | 3 |
| $rs$ | uniform | 0 | 3 | 0 | 3 |
| $\delta_s$ | uniform | 0 | 3 | 0 | 3 |
| $\epsilon_{pool}$ | uniform | 5 | 250 | 5 | 250 |
| $\epsilon_{seq}$ | uniform | 0.0001 | 0.01 | 0.0001 | 0.01 |
| $m_{cw}$ | uniform | $10^{-13}$ | $10^{-3}$ | $10^{-13}$ | $10^{-3}$ |
| $m_{wc}$ | uniform | $10^{-13}$ | $10^{-3}$ | $10^{-13}$ | $10^{-3}$ |
| $m_{cc}$ | uniform | $10^{-13}$ | $10^{-3}$ | $10^{-16}$ | $10^{-4}$ |
| $m_{ww}$ | uniform | $10^{-13}$ | $10^{-3}$ | $10^{-16}$ | $10^{-4}$ |
| $m_{aa}$ | uniform | 0 | $10^{-8}$ | $10^{-16}$ | $10^{-4}$ |
| $P_{cw}$ | beta | 0 | 0.2 | 0 | 0.2 |
| $P_{wc}$ | beta | 0 | 0.2 | 0 | 0.2 |
| $P_{no}$ | beta | 0 | 0.1 | 0 | 0.1 |

We chose a set of statistics (Table S1) to capture the patterns of relative diversity and differen-tiation among populations (Fraïsse et al., 2021; Jay, Boitard, & Austerlitz, 2019). These statistics were computed exclusively for polymorphic sites across all populations. Specifically, we considered the following statistics: (i) expected heterozygosity, both within each population and between all pairs of populations (Nei & Roychoudhury, 1974); (ii) Pairwise *F_ST_* values, computed for all possible population pairs (Bhatia, Patterson, Sankararaman, & Price, 2013); (iii) proportion of SNPs exhibiting fixed differences between populations (Fraïsse et al., 2021); (iv) proportion of SNPs exclusive to each individual population (Fraïsse et al., 2021) and (V) various D-statistics using different combinations of the P1, P2, and P3 populations (adapted from Malinsky, Matschiner, & Svardal, 2021). To capture the variation across loci, we considered the mean and standard deviation of the aforementioned statistics. We also considered the 5% and the 95% quantiles of *F_ST_* because they are expected to provide insights into the impact of barriers to gene flow. In total, we considered a set of 61 summary statistics for both scenarios of ecotype formation (Table S1). Notably, all of these summary statistics are relative measures of diversity and differentiation (e.g., *F_ST_*) and their values are contingent upon the relative branch lengths of coalescent trees.

Given that all summary statistics depend on the relative branch lengths of coalescent trees, we could increase the efficiency of our simulations by inferring relative demographic parameters scaled by the ancestral effective population size *N_re f_* . This approach allowed us to estimate relative effective sizes, such as *n*_1_ = *N*_1_*/N_re f_*, relative divergence times, such as *δ_s_*= Δ*_s_/*4*N_re f_*, and scaled migration rates, such as 4*N*_1_*m*_21_. It should be noted that relative parameters are denoted with lowercase letters (e.g., *n*_1_), while absolute parameters are represented with uppercase letters (e.g., *N*_1_). Scaled migration rates (forward in time) specify which population is receiving immigrants, indicated by the subscript next to *N*. We estimated relative parameters by performing coalescent simulations, with the ancestral effective population size set at *N_re f_* = 25000 and the mutation rate at *µ* = 1.5 × 10*^−^*^8^ per site per generation. These parameter values were chosen based on previous *L. saxatilis* studies (Butlin et al., 2014). In order to obtain absolute parameter estimates, we used a re-scaling factor defined as *f* = *obs*[*S*]*/E*[*S*]. This factor depends on the observed number of SNPs (*obs*[*S*]) and on the expected number of SNPs calculated using the parameter estimates of a given scenario (*E*[*S*]). Under the assumptions of the infinite sites mutation model, the expected number of segregating sites (*E*[*S*]) was determined by considering the expected total branch length (*E*[*T*]), the mutation rate per site (*µ*), and the number of sites (*S*). This relationship can be expressed as *E*[*S*] = *E*[*T*] · *µ* · *S* (Hudson, 1990). To obtain *E*[*T*] we simulated 100,000 gene trees, using the parameter estimates from the ecotype formation scenario that received the highest posterior support. The absolute effective population sizes and times of events in generations were then obtained by multiplying the rescaling factor with the corresponding relative values, i.e., *N_e_* = *f ×n_e_* and *T_s_* = *f ×t_s_*, respectively.

### Simulation study

To mitigate the computational load associated with simulating entire genomes, we instead simulated sets of *L* loci. We assumed complete independence and allowed for free recombination between all pairs of loci (*r_b_* = 0.5). Additionally, within each locus, we assumed no recombination (*r_w_* = 0.0). We performed 5 × 10^5^ simulations for each scenario of ecotype formation. In each simulation, we generated *L* = 300 independent loci, each consisting of *b* = 2000 base pairs, sampling 100 diploid individuals from each population. Pool-seq data were generated for each population by assuming 20 pools, with each pool containing 5 individuals. We assumed that all loci within a subset (i.e., each set of *L* = 300 independent loci) share a common demographic history. However, we modeled a migration rate of zero for a proportion of loci, *P_no_*, *P_cw_* and *P_wc_*, to account for the influence of selection effects resulting from barrier loci. Most of the parameter values were randomly sampled from uniform or log-uniform prior distributions, as outlined in Table 1. The proportions of the genome without migration (*P_no_*, *P_cw_* and *P_wc_*) were sampled from a Beta distribution specifically designed to reflect a low proportion of loci without migration as a prior assumption. We truncated the distribution of these parameters, replacing values below the minimum prior range or above the maximum prior range with the corresponding minimum or maximum values (Table 1).

We performed a leave-one-out cross-validation (Csilléry, François, & Blum, 2012) to assess the accuracy of parameter estimation and model choice. Briefly, we randomly selected a simulation and used its summary statistics as pseudo-observed data. The remaining simulations were then utilized to infer the parameters associated with the selected simulation. This process was repeated for a total of *n* pseudo-observed datasets. In this context, we define accuracy as the proximity of a specific point estimate to the true parameter value. We computed the prediction error for parameter inferences as:

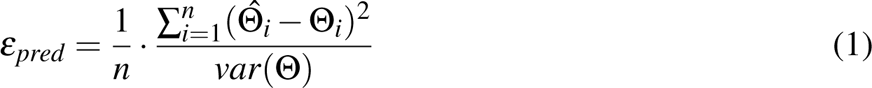

where Θ*_i_* represents the true parameter value of the *i^th^*pseudo-observed dataset, 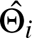 denotes the estimated parameter value, and *var*(Θ) corresponds to the variance of the true parameter values. We evaluated the prediction error for parameter inference with *n* = 5000 pseudo-observed datasets and considering three different point estimates (mode, median and mean of the posterior distribution, at two tolerance values (0.005 or 0.01). To evaluate the prediction error for model choice, we used *n* = 1000 pseudo-observed datasets. We used two posterior probability thresholds to define the estimated model for each pseudo-observed dataset. The first threshold was set at 0.5, assigning a dataset to the model with a posterior probability higher than 0.5. For a more stringent criterion, we used a threshold of 0.9, only assigning a dataset to a model if its posterior probability exceeded this threshold. If the posterior probability did not meet the 0.9 threshold, it was classified as ”unclear”.

### Pool-seq data from *Littorina saxatilis*

We contrasted scenarios of ecotype formation using previously published pool-seq data (Morales et al., 2018) from *L. saxatilis* populations. We compared populations sampled from Sweden (Arsklovet) and Spain (Burela). At each of those countries, 100 females from the Crab ecotype and another 100 females from the Wave ecotype were sequenced in separate pools (Morales et al., 2019). DNA was extracted from groups of individuals by combining foot muscle tissue samples from five snails in one tube. Subsequently, the reads obtained were trimmed using Trimmomatic v0.36 (Bolger, Lohse, & Usadel, 2014) and mapped against the reference genome of *L. saxatilis*, which was generated from one individual of the Crab ecotype (Westram et al., 2018). The mapping process was performed using CLC v5.0.3 from Qiagen Bioinformatics (www.qiagenbioinformatics.com). After mapping the reads, only those with a mapping score higher than Q20 were retained for further analysis. The resulting BAM files were processed using SAMtools v1.3.1 (Danecek et al., 2021), BEDtools v2.25.0 (Quinlan & Hall, 2010), and Picard tools v2.7.1. For each set of BAM files, reads with a base quality lower than 30 that mapped to contigs shorter than 500 base pairs were filtered out. To minimize potential artifacts, sites with a coverage lower than 14x or higher than 204x were excluded. This allowed us to discard low-coverage sites that lacked reads for most individuals (<14x), as well as sites in potentially repetitive or duplicated regions that led to unusually high coverage (>204x). Additionally, we excluded sites that had fewer than two minor-allele reads observed across all populations.

Recently, the significance of chromosomal inversions in the adaptive divergence of *L. saxatilis* has been highlighted (Faria et al., 2019; Koch et al., 2021; Morales et al., 2019). Each inversion is likely to have its own unique evolutionary history, which can be influenced by a range of demographic and selective processes, including divergent and balancing selection. It is important to note that the evolutionary dynamics of inversions may differ from the overall population history. Consequently, to ensure unbiased estimates, it would be necessary to use inversion-specific inference methods that take into account specific features, such as variable recombination rates between homozygotes and heterozygotes. As our primary goal was to infer the demographic history rather than the specific dynamics of inversions, we opted to remove regions that could potentially be associated or linked with the reported inversions (Westram et al., 2021). The list of retained and excluded contigs can be found in the Supplementary information. By removing these regions, we focused the analysis on genomic regions less likely to be influenced by inversion-specific processes, thus ensuring a more conservative inference of the neutral demographic history of those *L. saxatilis* populations. Due to uncertainty about the precise positions of breakpoints for many inversions we made the decision to exclude a total of 3671 contigs located within inversions or buffer regions. This accounts for approximately 3.3% of the entire genome pool-seq dataset. These excluded contigs were distributed across the genome, with roughly one-third of them located in chromosomes 10 and 12. Furthermore, we removed all contigs that did not map to known collinear regions (Westram et al., 2018). Consequently, around 80% of the remaining contigs were excluded from our analysis, leaving only the known collinear regions.

To reduce computational burden, we saved parameter and summary statistic tables from the simulation study, which were reused to perform model choice and estimate parameters for the *L. saxatilis* populations. We chose the scenario with the highest posterior probability as the preferred model and performed parameter inference for the selected scenario. Following our strategy of using subsets of loci, we treated each contig in the *L. saxatilis* dataset as an independent locus. To obtain posterior probabilities, we combined 1000 subsets, each consisting of 300 randomly selected loci (*L* = 300). The selection process for each subset involved randomly choosing 300 contigs without replacement. From these selected contigs, we then randomly extracted a window of *b* = 2000 base pairs. We computed summary statistics for each subset consisting of the 300 selected windows. Given the reduced number of mapped collinear contigs available (∼ 8000), contigs were reused in different subsets. Nevertheless, due to the high probability of selecting different 2000 base pairs windows, each subset likely represents a distinct combination of loci. The independent posterior samples obtained from the 1000 subsets of loci were merged, taking into account the distance between the mean summary statistics of each subset and the overall mean across all loci in the genome. To achieve this, we used the Epanechnikov kernel, which assigns greater weight to subsets of loci with means that are closer to the overall mean. This approach was designed to minimize the influence of outlier subsets of loci on the posterior estimates, given that demographic history is expected to affect all loci similarly across the genome. All steps were performed using the R package *poolABC* (Carvalho et al., 2023). Here we report the mean of the merged posterior distribution as a point estimate and the 95% credible intervals using weighted quantiles. To rescale the relative parameters, we calculated the number of SNPs per window, considering that all remaining sites were monomorphic. We converted the time of events from generations to years by assuming a generation time of 0.5 years (Butlin et al., 2014).

## RESULTS

### Accuracy of ABC point estimates

Results from the simulation study showed that prediction errors were lower, for all parameters, when using the mean or median with the regression-based adjustment (supplementary Tables S2 and S3). Thus, unless specified, hereafter we summarize results obtained using the regression-based adjustment and the mean as a point estimate, with a tolerance of 0.01.

For the relative effective sizes of present-day populations (Figure 1), the prediction errors ranged between 0.190 and 0.193 for the single origin scenario, and between 0.119 and 0.135 for the parallel origin scenario. These results indicate that the means of the posterior distributions provide accurate point estimates of these parameters. When considering the relative sizes of the ancestral populations (absolute values indicated by *NA*_1_ and *NA*_2_ in Figure 1), the prediction errors were similar in the two scenarios of ecotype formation (Table 2). In both models, the prediction error for the relative sizes of ancestral populations, *na*_1_ and *na*_2_, was higher (ranging from 0.863 to 0.911) than the error observed for the present-day populations.

**Table 2:** Prediction errors for relative parameters estimation. Prediction errors were computed using the mean of the posterior distribution, obtained after the regression adjustment and a tolerance of 0.01. *n*_1_ to *n*_4_ - relative population sizes of the extant populations; *na*_1_ and *na*_2_ - relative population sizes of the ancestral populations; *rs*_1_ and *rs*_2_ - relative time of the recent split event; *δ_s_* - relative time interval between *rs* and the ancient split event (*t_As_*); *ε_pool_*- experimental error introduced by the pooling procedures; *ε_seq_* - error associated with sequencing and mapping errors; *m_cw_*_1_*, m_cw_*_2_ - probability per generation that an individual migrates from the *N*_1_ or *N*_3_ (Crab) population to the *N*_2_ or *N*_4_ (Wave) population (forward in time), *m_wc_*_1_*, m_wc_*_2_ - probability per generation that an individual migrates from the *N*_2_ or *N*_4_ (Wave) population to the *N*_1_ or *N*_3_ (Crab) population (forward in time); *m_cc_* - probability per generation that an individual migrates from one Crab population to the other (forward in time); *m_ww_* - probability per generation that an individual migrates from one Wave population to the other (forward in time); *m_aa_*- probability per generation that an individual migrates from one ancestral population to the other (forward in time); 4*N_i_m_cw_*_1_ and 4*N_i_m_wc_*_1_ - average number of immigrants per generation (4*Nm*) from Crab to Wave and from Wave to Crab (respectively) in the first location; 4*N*_4_*m_cw_*_2_ and 4*N*_3_*m_wc_*_2_ - equivalent immigration rates at the second site; 4*N_i_m_cc_*_1_ and 4*N_i_m_cc_*_2_ - average number of immigrants per generation from the Crab population in the second location to the first and vice-versa (respectively); 4*N_i_m_ww_*_1_ and 4*N_i_m_ww_*_2_ - average number of immigrants per generation from the Wave population in the second location to the first and vice-versa (respectively); 4*NA*_1_*m_aa_*_1_ and 4*NA*_2_*m_aa_*_2_ - average number of immigrants per generation from *na*_2_ to *na*_1_ and vice-versa (respectively); *P_cw_*- proportion of the simulated loci where no migration occurs from the Crab to the Wave population; *P_wc_* - proportion of the simulated loci where no migration occurs from the Wave to the Crab population and *P_no_* - proportion of the simulated loci where no migration occurs between ecotypes.

| parameter | single | parallel | parameter | single | parallel |
| --- | --- | --- | --- | --- | --- |
| $n_1$ | 0.182 | 0.119 | $m_{ww}$ | 0.196 | 0.148 |
| $n_2$ | 0.182 | 0.132 | $m_{aa}$ | 1.021 | 0.977 |
| $n_3$ | 0.193 | 0.135 | $4N_im_{cw1}$ | 0.330 | 0.287 |
| $n_4$ | 0.190 | 0.129 | $4N_im_{wc1}$ | 0.319 | 0.266 |
| $na_1$ | 0.863 | 0.883 | $4N_im_{cw2}$ | 0.333 | 0.308 |
| $na_2$ | 0.911 | 0.907 | $4N_im_{wc2}$ | 0.332 | 0.326 |
| $rs_1$ | 0.616 | 0.505 | $4N_im_{cc1}$ | 0.254 | 0.143 |
| $rs_2$ | 0.631 | 0.513 | $4N_im_{cc2}$ | 0.263 | 0.146 |
| $\delta_s$ | 0.547 | 0.992 | $4N_im_{ww1}$ | 0.266 | 0.133 |
| $\epsilon_{pool}$ | 0.082 | 0.138 | $4N_im_{ww2}$ | 0.252 | 0.140 |
| $\epsilon_{seq}$ | 0.021 | 0.022 | $4NA_1m_{aa1}$ | 0.934 | 0.979 |
| $m_{cw1}$ | 0.469 | 0.449 | $4NA_2m_{aa2}$ | 0.952 | 0.981 |
| $m_{cw2}$ | 0.472 | 0.465 | $P_{cw}$ | 0.800 | 0.827 |
| $m_{wc1}$ | 0.451 | 0.427 | $P_{wc}$ | 0.753 | 0.814 |
| $m_{wc2}$ | 0.468 | 0.480 | $P_{no}$ | 0.094 | 0.140 |
| $m_{cc}$ | 0.211 | 0.154 | | | |

This suggests that point estimates are much less accurate for the ancestral effective sizes compared to the present-day populations (Figure 2 and supplementary Figures S1 and S2). The prediction errors for the relative time of the recent split event between populations was lower in the parallel scenario than in the single scenario. Specifically, at the first location (*rs*_1_), the prediction error was 0.505 for the parallel scenario and 0.616 for the single scenario. Likewise, at the second location (*rs*_2_), the prediction error was 0.513 for the parallel scenario and 0.631 for the single scenario. In contrast, the prediction error for the relative time interval between split events (*δ_s_*) was lower for the single origin scenario (0.547) compared to the parallel scenario (0.992).

**Figure 2:**
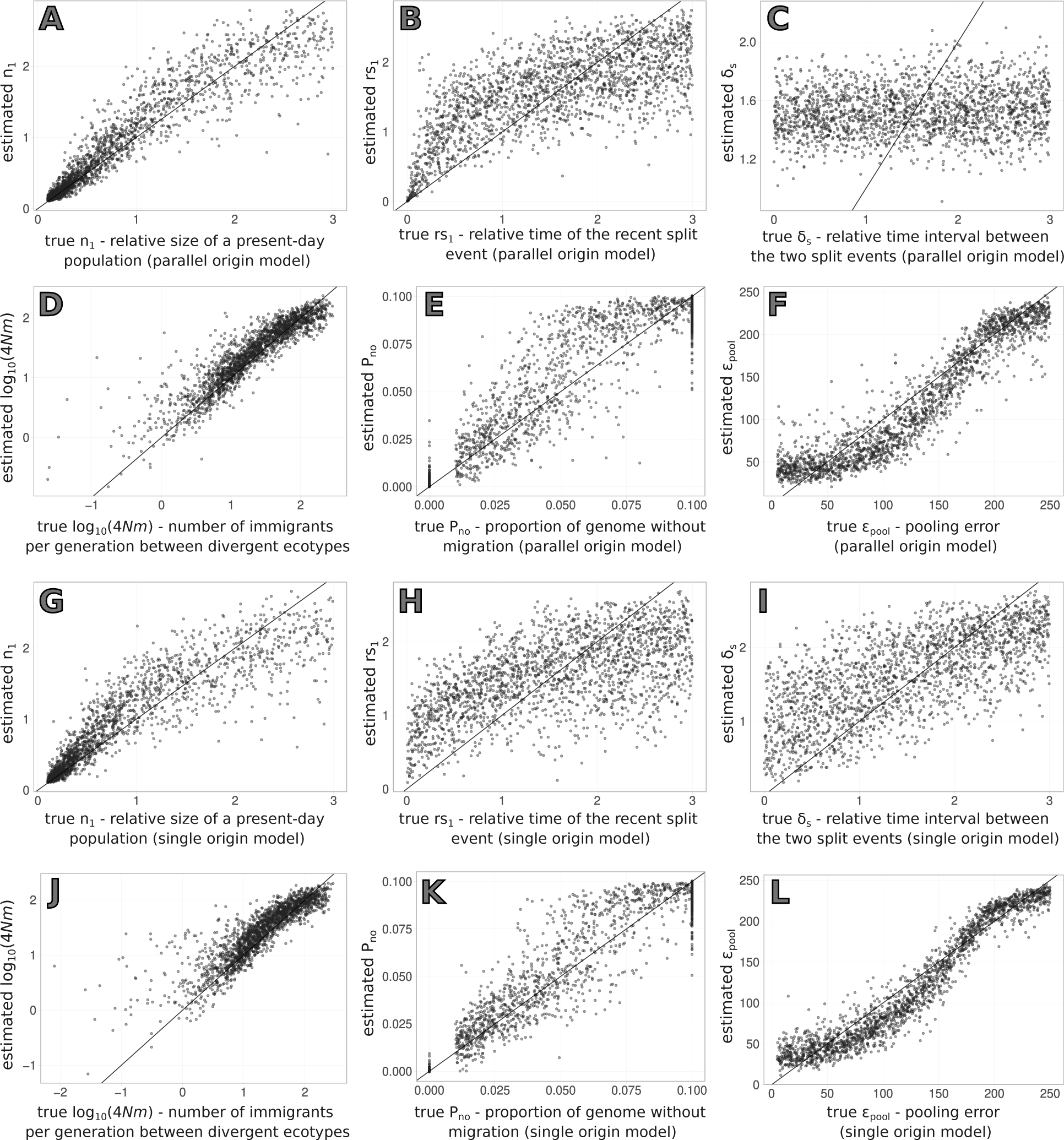
Results of the cross-validation for parameter estimation. The y-axis displays the estimated values, plotted against the true parameter values on the x-axis. Estimates correspond to the mean of the posterior obtained with a tolerance rate of 0.01. The top 6 panels (A-F) show the results for the parallel origin scenario, while the bottom 6 panels (G-L) correspond to the single origin scenario. Parameters shown here are: A and G - relative size of a present-day population (*n*_1_); B and H - relative time of the recent split event in the first location (*rs*_1_); C and I - relative time interval between the two split events (*δ_s_*); D and J - average number of immigrants per generation in *log*_10_ scale; E and K - proportion of the simulated loci where no migration occurs between ecotypes (*P_no_*); F and L - pooling error.

Although we parameterized our models using prior immigration rates *m_i j_*, which represent the probability of a lineage migrating from population *i* to *j* in each generation, we focus here on the average number of immigrants per generation denoted as 4*N_j_m_i j_*. In this context, *N_j_*represents the effective size of the population that receives immigrants. This measure takes into consideration both migration, which is proportional to *m_i j_*, and genetic drift, which is proportional to *N_j_*. Notably, when 4*N_j_m_i j_* > 1, migration occurs at a higher rate than drift. Prediction errors for average number of immigrants per generation between the divergent ecotypes were similar for the two scenarios of ecotype formation, ranging from 0.266 to 0.333 (Table 2).

Regarding the number of immigrants between populations of the same ecotype inhabiting different locations, the prediction errors were slightly lower in the parallel scenario (∼ 0.150) compared to the single origin scenario (∼ 0.200). The prediction errors for the number of immigrants between ancestral populations were high in both scenarios, ranging from 0.934 to 0.981. In contrast to the high prediction error observed for the proportion of loci without migration from the Crab to the Wave ecotype (*P_cw_*) and vice versa (*P_wc_*), which was approximately 0.8 in both scenarios, the proportion of loci without migration in either direction (*P_no_*) was accurately estimated. The prediction error for *P_no_* was 0.140 in the parallel scenario and 0.094 in the single origin scenario (Table 2).

Finally, and although our primary goal was to infer demographic parameters while explicitly modeling pool-seq data and treating pooling and sequencing errors as nuisance parameters, we also provide the prediction error for these parameters. The accuracy of the inference of the pooling error was similar to that of other parameters, with errors ranging from 0.082 to 0.138 (Table 2). This parameter was reasonably well estimated by the posterior mean when simulations were done with pooling errors above 150% (Figure 2). For the sequencing error, prediction error was low (∼ 0.02, Table 2 and supplementary Figures S1 and S1), probably because of the number of individuals considered here.

### Accuracy of model choice

The results of our simulation study demonstrate that the combination of ABC with pool-seq data effectively enables to distinguish between the two scenarios of ecotype formation considered here. Using a 50% posterior probability threshold, the correct model was successfully inferred for 988 out of 1,000 pseudo-observed datasets simulated under the parallel origin scenario, with a mean posterior probability of 0.976. Similarly, for the single origin scenario, the correct model was inferred for 989 pseudo-observed datasets, with a mean posterior probability of 0.995 (Figure 3A). In cases where the model with the highest posterior probability was incorrect, its posterior probability was lower. Specifically, when the parallel origin scenario was incorrectly inferred as single origin, the posterior probability was 0.671 and, when the single origin scenario was mistakenly inferred as parallel origin, the posterior probability was 0.899. Even with a more stringent threshold of 90% posterior probability, ABC analysis continued to successfully differentiate between the two scenarios. Out of the pseudo-observed datasets analyzed, the correct model was inferred for 913 datasets of parallel origin (with 87 classified as unclear), and for 973 datasets simulated under single origin, with 6 incorrectly assigned to parallel and 21 classified as unclear (Figure 3B).

**Figure 3:**
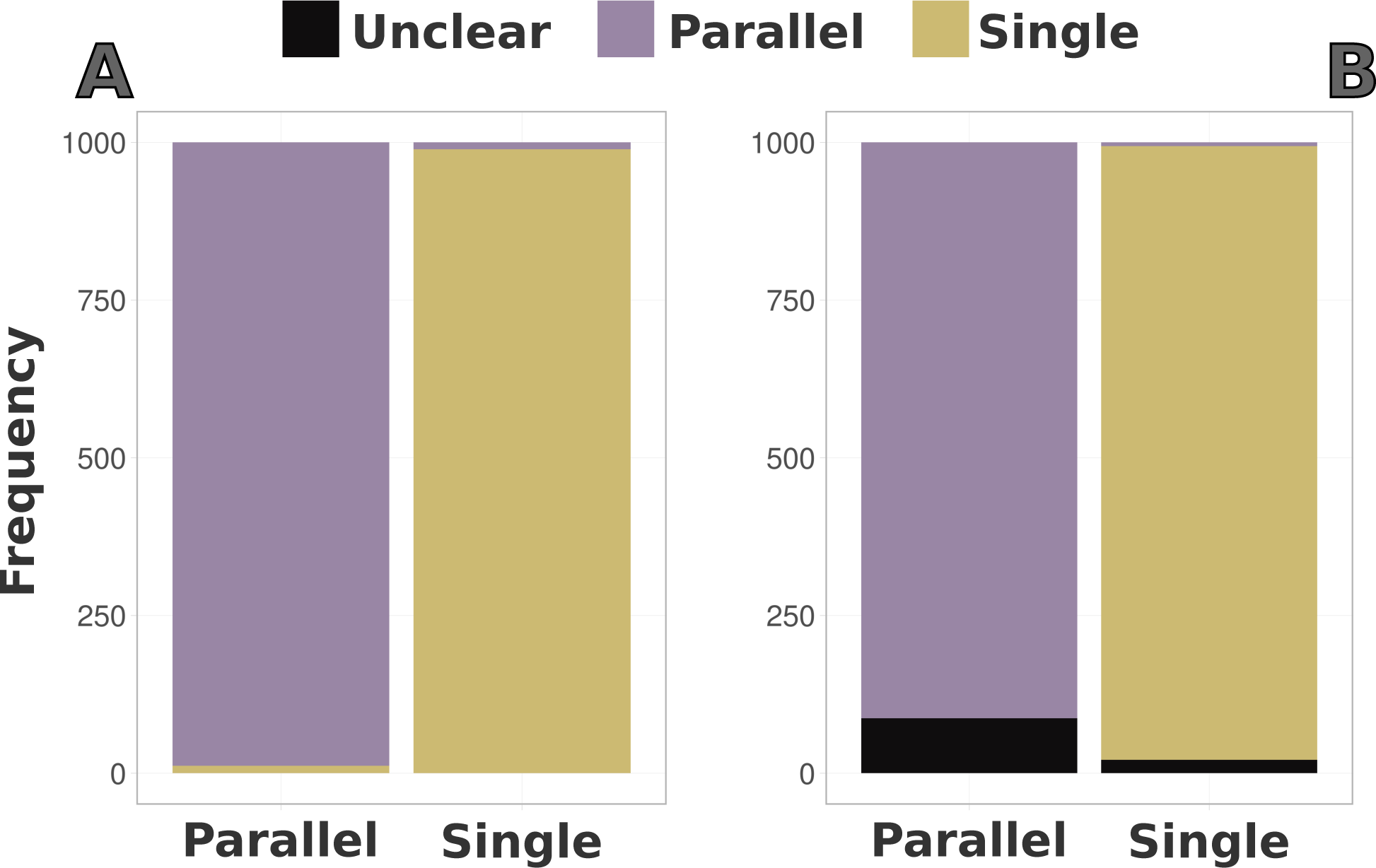
Model misclassification for the parallel and single origin models. Misclassification is based on the confusion matrix obtained using two different thresholds: (A) Simulations assigned to a model if posterior probability was above 0.5 or (B) Simulations assigned to a model only if posterior probability was above 0.9 (B).

### Model choice and parameter inference of *Littorina saxatilis*

Our analysis of the Crab and Wave ecotypes sampled from two geographically distant locations (Arsklovet in Sweden and Burela in Spain) provides strong support for the parallel origin model. The posterior probability obtained using the rejection algorithm was 0.993, while the posterior probability obtained using logistic regression was 1.000. Thus, we report here the parameter estimates obtained using the parallel origin scenario. To simplify the presentation of results, we re-scaled relative parameters to absolute effective sizes and time of events in years, using *k* to indicate thousands (Table 3, but refer to Table S4 for the relative estimates). The re-scaling process was conducted after combining the posterior distributions from 1000 subsets of loci, assigning more weight to subsets of loci with summary statistics closer to the overall mean of the entire genome.

**Table 3:** Absolute parameter estimates for *Littorina saxatilis* populations. Results are shown for the parallel origin scenario using the Arsklovet and Burela populations. For this model *N*_1_ and *N*_2_ correspond, respectively, to the absolute size of the Arsklovet Crab and Wave populations, while *N*_3_ and *N*_4_ correspond to the absolute size of the Burela Crab and Wave populations, respectively. For each parameter, the value outside brackets corresponds to the re-scaled mean of the posterior distribution and in-between brackets is the 95% credible interval. *RS*_1_, *RS*_2_ and Δ*_s_*are presented in years. Parameters indicated here are the same as in table 2, except for *P_no_*, which is converted to the percentage of the genome where no migration occurs between ecotypes.

| parameter | absolute estimate |
| --- | --- |
| $N_1$ | 8842 (6434 - 16442) |
| $N_2$ | 11562 (7404 - 23233) |
| $N_3$ | 36816 (17565 - 88184) |
| $N_4$ | 25573 (12204 - 59078) |
| $NA_1$ | 42654 (7098 - 138326) |
| $NA_2$ | 59906 (8068 - 142922) |
| $RS_1$ | 14675 (2655 - 115400) |
| $RS_2$ | 57296 (10723 - 241318) |
| $\Delta_s$ | 220941 (31604 - 298711) |
| $4N_2m_{cw1}$ | 15.8 (4.2 - 35.4) |
| $4N_1m_{wc1}$ | 10.4 (2.8 - 23.3) |
| $4N_4m_{cw2}$ | 8.3 (1.8 - 20.4) |
| $4N_3m_{wc2}$ | 23.4 (4.7 - 62.4) |
| $4N_1m_{cc1}$ | 0.5 (0.2 - 1.0) |
| $4N_3m_{cc2}$ | 2.3 (0.8 - 4.7) |
| $4N_2m_{ww1}$ | 0.3 (0.1 - 0.6) |
| $4N_4m_{ww2}$ | 0.7 (0.2 - 1.4) |
| $4NA_1m_{aa1}$ | 3.3 (0.2 - 12.9) |
| $4NA_2m_{aa2}$ | 4.2 (0.2 - 15.5) |
| $P_{no}$ | 3.7 (0.5 - 8.8) |

Estimates based on the parallel origin scenario show that, in Spain, the present-day Crab population has a larger effective size of approximately 37k (95% CI: 18k - 89k) compared to the Wave population, which has an effective size of approximately 26k (95% CI: 12k - 59k). In contrast, the present-day populations in Sweden exhibit smaller effective sizes relative to the Spanish populations (Table 3), with the Wave population having a slightly larger effective size of approximately 12k (95% CI: 7k - 23k) compared to the Crab population, which has an effective size of ∼ 9k (95% CI: 6k - 16k). Our parameter estimates suggest that the two ecotypes diverged approximately 15,000 years ago (95% CI: 3k to 115k years) in Sweden (*RS*_1_), with a much older split between Crab and Wave ecotype populations in Spain (*RS*_2_), occurring ∼ 57,000 years ago (95% CI: 11k to 241k years). This divergence process was accompanied by gene flow in both countries. However, there were differences in the pattern, with higher scaled immigration (4*Nm*) from the Crab into the Wave ecotype in Sweden and from the Wave into the Crab ecotype in Spain (Table 3). Parameter estimates suggest that the separation between Spanish and Swedish populations (*T_As_*) occurred 278k years ago (95% CI: 42k to 540k years). The point estimates also supported a larger ancestral effective size of the Spanish population (mean ∼ 60k, 95% CI: 8k - 142k) compared with the Swedish population (mean ∼ 43k, 95% CI: 7k - 138k). Lastly, we inferred a mean proportion of loci without migration in either direction (*P_no_*) of approximately 4% with an upper CI close to 9% (Table 3). Furthermore, the proportion of loci without migration from Crab to Wave was approximately 2% (95% CI: 0% - 17.2%), and from Wave to Crab, it was around 3% (95% CI: 0% - 18.6%).

## DISCUSSION

We contrasted two possible scenarios of ecotype formation in *L. saxatilis* by using whole-genome data obtained with pool-sequencing. We used a recently developed model-based method that is specifically designed for analysing pooled-sequencing data (Carvalho et al., 2023). This method combines an ABC (Approximate Bayesian Computation) inference framework with the explicit modeling of the various sources of error associated with pool-sequencing. By incorporating those sources of error into the analysis, we aimed to avoid biases in demographic estimates due to poolseq errors.

### Pool-seq differentiates between complex scenarios of ecotype formation

The prediction errors obtained with the simulation study show that, for the datasets analysed here, the means of the posterior distributions provide accurate point estimates for most demographic history parameters of both scenarios of ecotype formation. The clear exceptions were the parameters related with the ancestral populations, such as the relative sizes of those populations and the migration between them. The high uncertainty associated with these parameters suggests that the summary statistics used do not contain information about older events. This observation is supported by the posterior distributions of these parameters in *L. saxatilis*, as they show minimal deviation from their prior distributions (Figure 4). Most of the remaining demographic history parameters exhibited similar prediction errors for both scenarios (Table 2). However, the prediction error for the relative time interval between split events (*δ_s_*) is higher for the parallel origin than the single origin scenario. This disparity can be explained by the underlying topology of the parallel origin scenario. In this scenario, the diverging ecotypes found at each location have a shared evolutionary history and experience ongoing gene flow between them. Consequently, accurately inferring past events in such a complex context can prove challenging. Another possible explanation is the existence of a lower migration rate between ancestral populations in the single origin scenario (Table 2). Furthermore, there is also a slightly larger prediction error for migration rates between the same ecotypes in the single origin scenario. This can be attributed to the challenge of distinguishing between gene flow and incomplete lineage sorting within populations that share a common evolutionary history in the single origin model.

**Figure 4:**
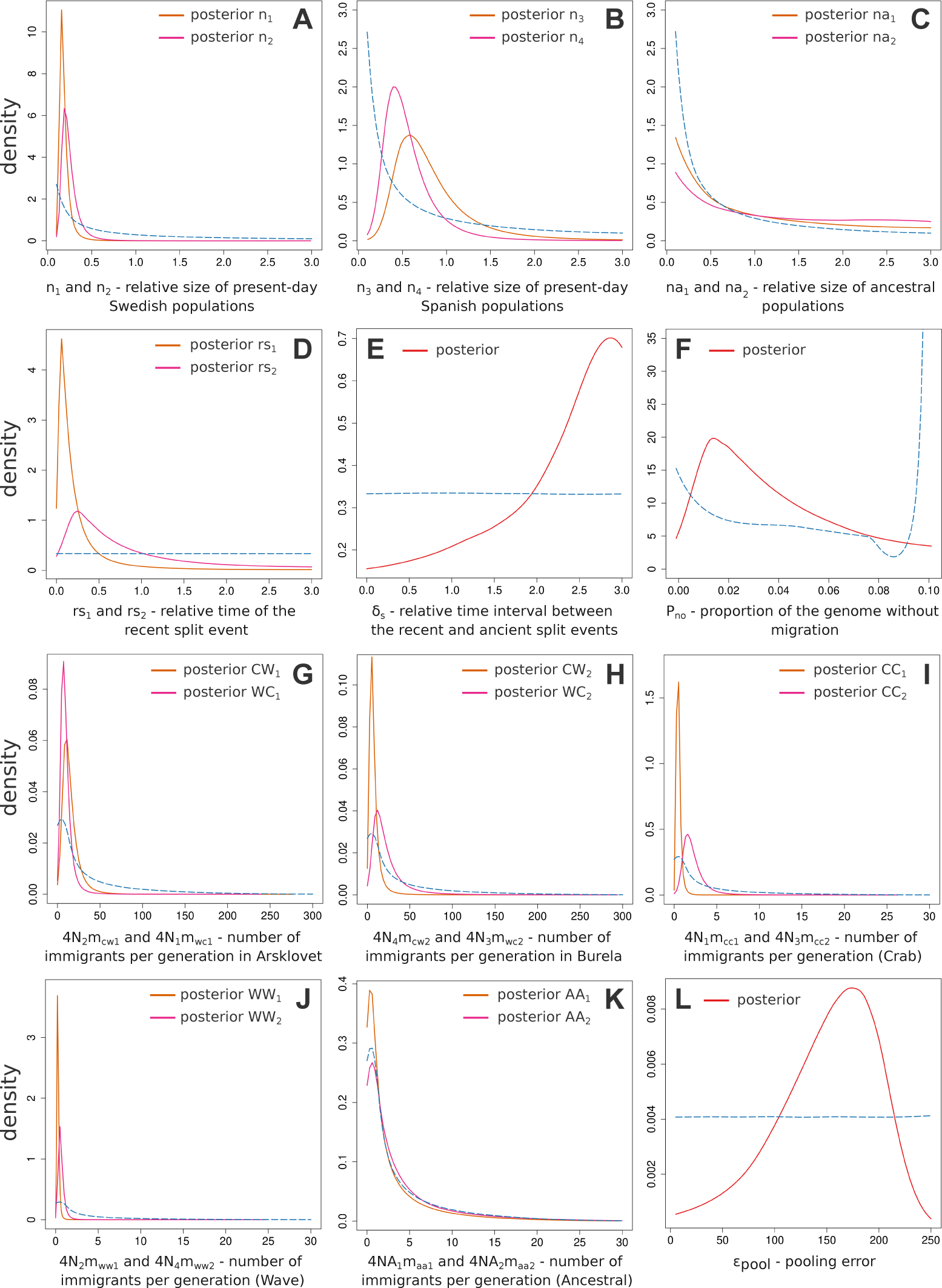
Posterior distributions of relative *L. saxatilis* parameters with regression adjustment and a tolerance of 0.01. Prior distributions are shown as a dotted blue line for reference. A - relative size of Arsklovet Crab (*n*_1_) and Wave populations (*n*_2_), B - relative size of Burela Crab (*n*_3_) and Wave populations (*n*_4_), C - relative size of ancestral populations (*na*_1_ and *na*_2_), D - relative time of the recent split events (*rs*_1_ and *rs*_2_), E - relative time interval between the two split events (*δ_s_*), F - proportion of the genome without migration (*P_no_*), G - average number of immigrants per generation (4*N*_2_*m_cw_*_1_ and 4*N*_1_*m_wc_*_1_) in Arsklovet, H - average number of immigrants per generation (4*N*_4_*m_cw_*_2_ and 4*N*_3_*m_wc_*_2_) in Burela, I - average number of immigrants per generation (4*N*_1_*m_cc_*_1_ and 4*N*_3_*m_cc_*_2_) between Crab populations, J - average number of immigrants per generation (4*N*_2_*m_ww_*_1_ and 4*N*_4_*m_ww_*_2_) between Wave populations, K - average number of immigrants per generation (4*NA*_1_*m_aa_*_1_ and 4*NA*_2_*m_aa_*_2_) between ancestral populations and L - pooling error.

Even though we discarded the majority of available contigs, results from our simulation study confirm that pool-seq provides sufficient information to differentiate between complex scenarios of ecotype formation. Remarkably, this distinction remains possible even when setting a high threshold for posterior probability, as indicated by the proportion of correctly assigned simulations with 90% posterior probability above 0.9 for both models (Figure 3B). This result is surprising considering the high connectivity between populations in our model, where virtually all pairs of populations have the potential for migration (Figure 1). Nevertheless, the single and parallel origin scenarios considered here exhibit different mean values for various summary statistics (Figure S3), which accounts for our ability to distinguish between them (Marin, Pillai, Robert, & Rousseau, 2014). Using this approach, we found evidence that supports a parallel origin of the Crab and Wave ecotypes, without allopatric separation, and after colonization of the different regions by an ancestral population, which is consistent with the findings of Butlin et al. (2014). However, we did not specifically include periods of allopatry (i.e. isolation without gene flow) in our models. Thus, it it remains a possibility that periods of allopatry played a role in the evolution of *L. saxatilis* ecotypes. Additionally, it is worth noting that the support for our findings is based on neutral loci and thus, it is still uncertain whether the alleles responsible for adaptive traits, potentially associated with inversions, have evolved in parallel.

### Recent parallel origin of *Littorina saxatilis* ecotypes

Our results support a relatively recent divergence of the ecotypes in both countries. Notably, the divergence in Sweden appears to be more recent, around 15,000 years ago, while in Spain, it took place approximately 57,000 years ago. These results are aligned with the hypothesis of a recent postglacial colonization of Swedish islands (Panova et al., 2011) and match previous estimates for Swedish populations (Carvalho et al., 2023). The estimates obtained by Butlin et al. (2014) date the divergence of the ecotypes to approximately 19k or 30k years ago, depending on the populations pairs analysed.

These estimates fall in-between the range of our estimates, which could be attributed to the fact that Butlin et al. (2014) included a single ecotype formation split event, thereby not accounting for potential variations between regions. The separation of ecotypes occurred relatively recently compared to the separation of populations in different regions, which we estimated to have taken place around 278,000 years ago. This suggests that the time interval between ancestral geographic structuring in Europe and the formation of the ecotypes was roughly 221k years in Spain and 263k years in Sweden. Nevertheless, it is likely that the split between different regions inferred here reflects an older split between Iberia and a northern refuge located outside of Sweden. Indeed, the formation of the Crab ecotype in Sweden required the arrival of predators selecting for the Crab ecotype, which occurred progressively in warming regions after the last glacial maximum. Thus, it is likely that *L. saxatilis* populations did not exist in Sweden for more than 200k years without ecotype formation. It is possible that the ecotypes were repeatedly formed and lost due to habitats changes associated with glacial fluctuations. This phenomenon could have had a more pronounced impact in Sweden, where such oscillations are expected to be more intense and frequent. Therefore, our estimates of approximately 15k years ago in Sweden and 57k years ago in Spain, may just reflect the most recent split between the two ecotypes.

Additionally, the time period until ecotype formation could have been influenced by local extinctions triggered by factors such as toxic algal blooms (Johannesson & Johannesson, 1995), which could lead to population reestablishment through individuals carrying alleles from source populations of both ecotypes (Butlin et al., 2014). We found that the effective size of the Crab ecotype population in Spain was larger than that of the Wave ecotype population, whereas the Wave population in Sweden displayed a slightly larger effective size compared to the Crab population. Our estimates also support a lower effective size for current populations compared to ancestral populations, confirming a lack of support for past population expansions (Butlin et al., 2014).

The larger effective sizes of both present-day and ancestral populations in Spain could be attributed to various factors. One possibility is that these differences could arise from variations in carrying capacities or population densities between the two geographical locations. Alternatively, this difference might reflect historical demographic dynamics, such as population bottlenecks caused by toxic algal blooms in Sweden (Johannesson & Johannesson, 1995), potentially leading to more pronounced founder events in Sweden compared to Spain. Furthermore, the observed differences could be attributed to the proximity of Spanish populations to glacial refugia (Blakeslee et al., 2021; Bosso et al., 2022), which could have acted as safe havens during past glacial periods, allowing populations in Spain to retain larger sizes. Additionally, the larger effective size of the Spanish Crab population could be due to patterns of gene flow, given that we estimated a higher migration rate from the Wave to the Crab population in Spain (Table 3). On the whole, our estimates suggest that *L. saxatilis* effective population sizes remained consistently large, possibly due to gene flow and a high degree of cold-tolerance.

As mentioned, we did not specifically include periods without gene flow in our models and thus, caution is needed when considering the extent of gene flow between the ecotypes during the divergence process. Nevertheless, our results indicate that the divergence process between ecotypes in both Spain and Sweden was accompanied by gene flow. Specifically, we observed high migration rates between the divergent ecotypes, with values exceeding 4*Nm* > 10 in most of the comparisons examined. Interestingly, we inferred a slightly higher migration rate from Crab to Wave ecotype in Sweden. This is consistent with the suggested higher net dispersal from the Crab to Wave ecotype as an explanation for the observed shift in cline centers towards the Wave habitat on Swedish islands (Westram et al., 2021).

In summary, our results argue in favor of a demographic history characterized by the spatial division of an ancestral population into a Spanish ancestral population and another ancestral population from which the ecotypes in Spain and Sweden, respectively, originated. Over time, distinct habitat-associated populations evolved at each of those geographic locations, while still experiencing gene flow between the diverging ecotypes. Thus, the available evidence strongly supports the conclusion that the Crab and Wave ecotypes observed in *L. saxatilis* arose as a result of divergent selection, despite the ongoing exchange of genetic material between them. This interplay between divergent selection and gene flow likely played a crucial role in shaping the evolutionary history of the *L. saxatilis* ecotypes, influencing the evolution of chromosomal rearrangements such as inversions (Ortiz-Barrientos, Engelstädter, & Rieseberg, 2016). Understanding the evolutionary processes that led to the emergence of these ecotypes not only sheds light on the population dynamics of *L. saxatilis*, but it also provides valuable insights into the mechanisms of parallel ecological adaptation and speciation. Despite some limitations, our findings indicate that combining pool-seq with ABC is an effective approach for investigating parallel evolution across a wide geographical range. The approach detailed here should be applied to other *L. saxatilis* populations, including additional sampling locations in Sweden, Spain and the United Kingdom.

## Acknowledgements

We thank Hernán E. Morales for processing the *L. saxatilis* pool-seq data. This work was funded by the strategic project UIDB/00329/2020 granted to cE3c from the Portuguese National Science Foundation — Fundação para a Ciência e a Tecnologia (FCT). JC was supported by an FCT Ph.D. scholarship (PD/BD/128350/2017). RF is funded by a FCT CEEC (Fundação para a Ciência e a Tecnologia, Concurso Estímulo ao Emprego Científico) contract (2020.00275.CEECIND) and by a FCT research project (PTDC/BIA-EVL/1614/2021). RKB was funded by the European Research Council (ERC-2015-AdG-693030-BARRIERS). VCS was supported by Fundação Ciência e Tecnologia (CEECINST/00032/2018/CP1523/CT0008) and by the Human Frontier Science Program (RGY0081/2020). We thank the National Network for Advanced Computing (RNCA) and INCD (https://incd.pt/) for access and use of their computing infrastructure, funded by FCT to VCS (2021.09795.CPCA).

## Author Contributions

JC performed the simulation study and demographic inference of *L. saxatilis* data. JC and VCS designed the demographic models and analysed the data. JC wrote the manuscript together with VCS, with support from RF and RKB. RF, RKB and VCS supervised the project. All authors planned the study and provided critical feedback that helped shape the analysis and manuscript.

---

**Table S1:** Set of summary statistics considered. Different combinations of D-statistics were employed to examine the level of introgression occurring between the distinct ecotypes. For D-statistic 1, P1 represented the Wave population in the first location (*N*_2_), P2 stood for the Wave population in the second location (*N*_4_), and P3 denoted the Crab population at the first location (*N*_1_). For D-statistic 2, P1 again referred to the Wave population at the first location (*N*_2_), while P2 now represented the Crab population in the second location (*N*_3_), and P3 was the Crab population at the first location (*N*_1_). D-statistic 3 retained P1 as the Wave population at the first location (*N*_2_), designated P2 as the Crab population at the first location (*N*_1_), and P3 as the Wave population in the second location (*N*_4_). In all these combinations, P4 was assumed to be an outgroup fixed, at all sites, for the major allele. For the proportion of exclusive SNPs, we computed this per location i.e. checking if each site was segregating in one population but not in the other population inhabiting the same location and globally by computing the proportion of sites that were segregating in only one population and not in the other three. For the proportion of shared or fixed differences SNPs, we computed this for the two populations that inhabit the same location and globally by comparing each population with the other three.

| summary statistic | four-populations |
| --- | --- |
| mean heterozygosity | 4 values (1 per population) |
| SD heterozygosity | 4 values (1 per population) |
| mean heterozygosity between populations | 6 pairwise values |
| SD heterozygosity between populations | 6 pairwise values |
| pairwise $F_{ST}$ | 6 pairwise values |
| SD $F_{ST}$ | 6 pairwise values |
| 5%$F_{ST}$ | 6 pairwise values |
| 95%$F_{ST}$ | 6 pairwise values |
| proportion of fixed differences | 3 values |
| proportion of exclusive SNPs | 5 values |
| proportion of shared SNPs | 3 values |
| mean D-statistic 1 | 1 value |
| mean D-statistic 2 | 1 value |
| mean D-statistic 3 | 1 value |
| SD D-statistic 1 | 1 value |
| SD D-statistic 2 | 1 value |
| SD D-statistic 3 | 1 value |
| total | 61 |

**Table S2:** Prediction errors for the parallel origin parameters. Parameter inference was performed using a simple rejection or a regression adjustment using a local linear regression. For each method, values are presented for two different tolerance rates. *n*_1_ to *n*_4_ - relative population sizes of the extant populations, *na*_1_ and *na*_2_ - relative population sizes of the ancestral populations, *rs*_1_ and *rs*_2_ - relative time of the recent split events, *δ_s_* - relative time interval between *rs* and the ancient split event (*t_As_*), *ε_pool_* - experimental error introduced by the pooling procedures, *ε_seq_* - error associated with sequencing and mapping errors, *m_cw_*_1_ and *m_cw_*_2_ - probability per generation that an individual migrates from the Crab population to the Wave population in the first and second locations, respectively (forward in time), *m_wc_*_1_ and *m_wc_*_2_ - probability per generation that an individual migrates from the Wave population to the Crab population in the first and second locations, respectively (forward in time), *m_cc_* - probability per generation that an individual migrates from one Crab population to the other (forward in time), *m_ww_* - probability per generation that an individual migrates from one Wave population to the other (forward in time), *m_aa_* - probability per generation that an individual migrates from one ancestral population to the other (forward in time), *P_cw_* - proportion of the simulated loci where no migration occurs from the Crab to the Wave population; *P_wc_* - proportion of the simulated loci where no migration occurs from the Wave to the Crab population, *P_no_* - proportion of the simulated loci where no migration occurs between ecotypes, 4*N*_2_*m_cw_*_1_ and 4*N*_1_*m_wc_*_1_ - average number of immigrants per generation from Crab to Wave and from Wave to Crab (respectively) in the first location, 4*N*_4_*m_cw_*_2_ and 4*N*_3_*m_wc_*_2_ - average number of immigrants per generation from Crab to Wave and from Wave to Crab (respectively) in the second location, 4*N*_1_*m_cc_*_1_ and 4*N*_3_*m_cc_*_2_ - average number of immigrants per generation from the Crab population in the second location to the first and vice-versa (respectively), 4*N*_2_*m_ww_*_1_ and 4*N*_4_*m_ww_*_2_ - average number of immigrants per generation from the Wave population in the second location to the first and vice-versa (respectively), 4*NA*_1_*m_aa_*_1_ and 4*NA*_2_*m_aa_*_2_ - average number of immigrants per generation from *na*_2_ to *na*_1_ and vice-versa (respectively).

| parameter | REJECTION |  |  |  |  |  | REGRESSION |  |  |  |  |  |
| --- | --- | --- | --- | --- | --- | --- | --- | --- | --- | --- | --- | --- |
|  | tolerance of 0.005 |  |  | tolerance of 0.01 |  |  | tolerance of 0.005 |  |  | tolerance of 0.01 |  |  |
|  | mode | median | mean | mode | median | mean | mode | median | mean | mode | median | mean |
| $n_1$ | 0.94 | 0.49 | 0.44 | 1.04 | 0.55 | 0.49 | 0.12 | 0.12 | 0.13 | 0.13 | 0.12 | 0.13 |
| $n_2$ | 0.92 | 0.49 | 0.44 | 1.03 | 0.55 | 0.48 | 0.13 | 0.12 | 0.13 | 0.13 | 0.13 | 0.13 |
| $n_3$ | 0.97 | 0.50 | 0.46 | 1.05 | 0.53 | 0.48 | 0.14 | 0.14 | 0.14 | 0.13 | 0.13 | 0.13 |
| $n_4$ | 1.01 | 0.53 | 0.47 | 1.05 | 0.53 | 0.49 | 0.12 | 0.12 | 0.13 | 0.12 | 0.12 | 0.13 |
| $na_1$ | 2.00 | 1.13 | 0.96 | 1.97 | 1.13 | 0.98 | 1.96 | 0.97 | 0.90 | 1.91 | 0.99 | 0.93 |
| $na_2$ | 1.99 | 1.14 | 0.98 | 1.95 | 1.12 | 0.97 | 1.93 | 1.01 | 0.93 | 1.92 | 0.96 | 0.91 |
| $rs_1$ | 1.49 | 0.79 | 0.80 | 1.37 | 0.79 | 0.80 | 0.74 | 0.51 | 0.50 | 0.82 | 0.52 | 0.51 |
| $rs_2$ | 1.37 | 0.73 | 0.77 | 1.42 | 0.78 | 0.80 | 0.72 | 0.49 | 0.48 | 0.81 | 0.53 | 0.52 |
| $\delta_s$ | 2.81 | 0.99 | 0.99 | 3.04 | 1.00 | 1.00 | 2.66 | 1.05 | 1.00 | 2.79 | 1.02 | 0.99 |
| $\epsilon_{pool}$ | 0.44 | 0.34 | 0.39 | 0.50 | 0.39 | 0.44 | 0.13 | 0.13 | 0.13 | 0.14 | 0.14 | 0.14 |
| $\epsilon_{seq}$ | 0.25 | 0.21 | 0.26 | 0.27 | 0.27 | 0.33 | 0.03 | 0.02 | 0.02 | 0.03 | 0.03 | 0.02 |
| $m_{cw1}$ | 1.61 | 0.80 | 0.85 | 1.80 | 0.84 | 0.87 | 0.66 | 0.49 | 0.47 | 0.61 | 0.45 | 0.45 |
| $m_{cw2}$ | 1.68 | 0.81 | 0.85 | 1.70 | 0.84 | 0.88 | 0.61 | 0.44 | 0.44 | 0.63 | 0.47 | 0.46 |
| $m_{wc1}$ | 1.55 | 0.81 | 0.85 | 1.55 | 0.84 | 0.88 | 0.62 | 0.46 | 0.45 | 0.54 | 0.42 | 0.43 |
| $m_{wc2}$ | 1.74 | 0.81 | 0.85 | 1.83 | 0.86 | 0.89 | 0.55 | 0.41 | 0.41 | 0.65 | 0.48 | 0.48 |
| $m_{cc}$ | 0.86 | 0.54 | 0.57 | 0.95 | 0.54 | 0.59 | 0.16 | 0.15 | 0.15 | 0.17 | 0.16 | 0.15 |
| $m_{ww}$ | 0.83 | 0.51 | 0.55 | 0.92 | 0.56 | 0.60 | 0.15 | 0.14 | 0.14 | 0.16 | 0.15 | 0.15 |
| $m_{aa}$ | 2.73 | 0.99 | 0.99 | 2.79 | 0.98 | 0.98 | 2.50 | 1.05 | 0.99 | 2.66 | 1.02 | 0.98 |
| $P_{cw}$ | 2.81 | 1.01 | 0.97 | 2.85 | 1.04 | 0.98 | 2.18 | 0.86 | 0.78 | 2.28 | 0.92 | 0.83 |
| $P_{wc}$ | 2.75 | 1.05 | 0.97 | 2.81 | 1.04 | 0.98 | 2.12 | 0.87 | 0.78 | 2.33 | 0.91 | 0.81 |
| $P_{no}$ | 0.75 | 0.30 | 0.30 | 0.79 | 0.31 | 0.32 | 0.28 | 0.17 | 0.14 | 0.28 | 0.18 | 0.14 |
| $4N_2m_{cw1}$ | 1.16 | 0.76 | 0.65 | 1.24 | 0.82 | 0.68 | 0.33 | 0.30 | 0.29 | 0.34 | 0.30 | 0.29 |
| $4N_1m_{wc1}$ | 1.17 | 0.78 | 0.67 | 1.21 | 0.82 | 0.70 | 0.34 | 0.29 | 0.28 | 0.33 | 0.28 | 0.27 |
| $4N_4m_{cw2}$ | 1.21 | 0.80 | 0.68 | 1.26 | 0.83 | 0.71 | 0.32 | 0.28 | 0.28 | 0.36 | 0.31 | 0.31 |
| $4N_3m_{wc2}$ | 1.18 | 0.78 | 0.67 | 1.25 | 0.83 | 0.70 | 0.34 | 0.29 | 0.29 | 0.39 | 0.34 | 0.33 |
| $4N_1m_{cc1}$ | 0.93 | 0.62 | 0.52 | 0.99 | 0.67 | 0.54 | 0.14 | 0.14 | 0.14 | 0.15 | 0.14 | 0.14 |
| $4N_3m_{cc2}$ | 0.94 | 0.62 | 0.52 | 0.99 | 0.66 | 0.55 | 0.17 | 0.16 | 0.16 | 0.15 | 0.14 | 0.15 |
| $4N_2m_{ww1}$ | 0.91 | 0.61 | 0.51 | 0.98 | 0.66 | 0.54 | 0.15 | 0.14 | 0.14 | 0.14 | 0.13 | 0.13 |
| $4N_4m_{ww2}$ | 0.94 | 0.62 | 0.51 | 1.00 | 0.67 | 0.56 | 0.14 | 0.13 | 0.13 | 0.15 | 0.14 | 0.14 |
| $4NA_1m_{aa1}$ | 1.68 | 1.16 | 0.98 | 1.67 | 1.16 | 0.99 | 1.65 | 1.12 | 0.98 | 1.65 | 1.12 | 0.98 |
| $4NA_2m_{aa2}$ | 1.65 | 1.17 | 0.99 | 1.68 | 1.16 | 0.99 | 1.61 | 1.12 | 0.98 | 1.66 | 1.12 | 0.98 |

**Table S3:**
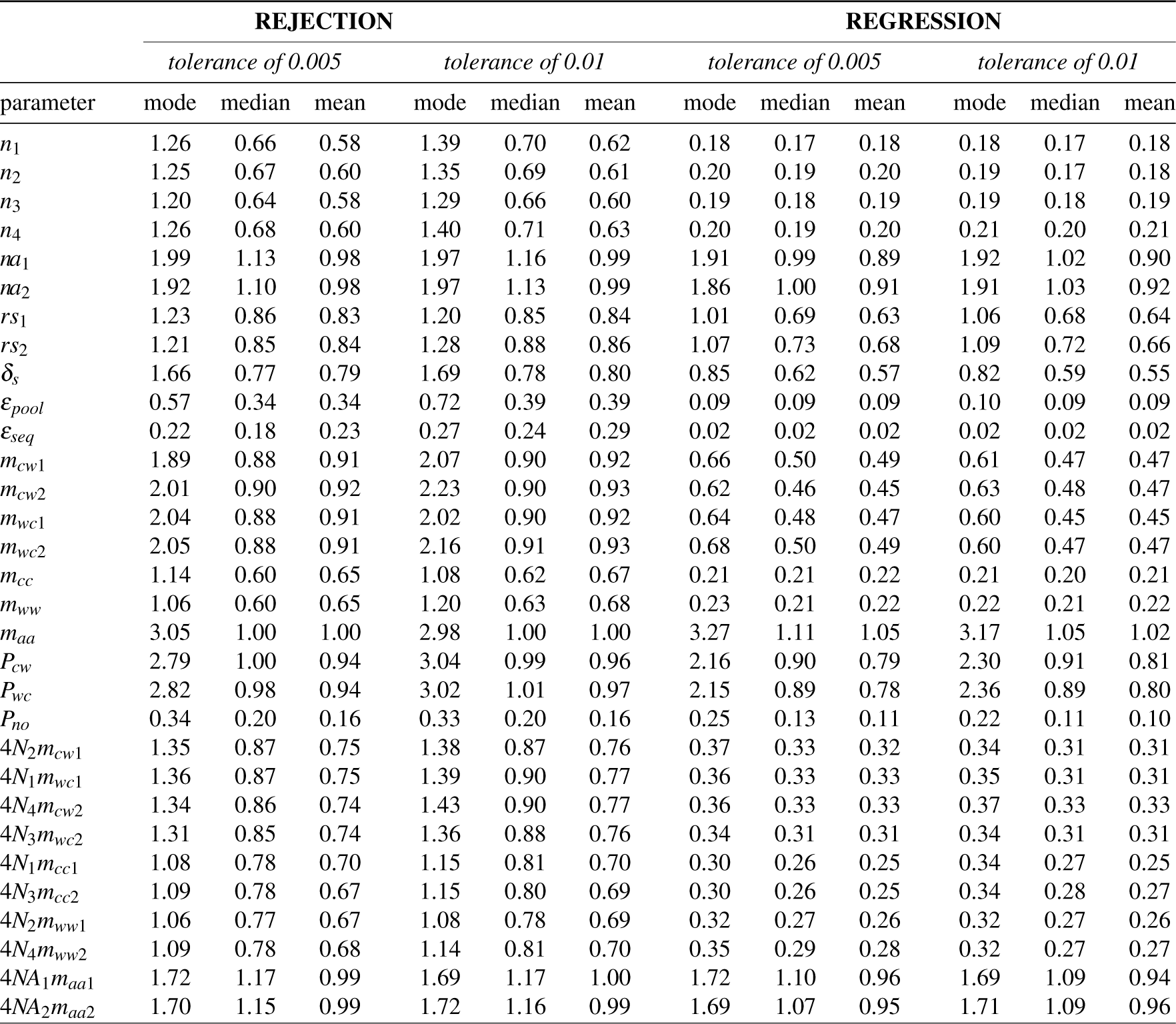
Prediction errors for the single origin parameters. Parameter inference was performed using a simple rejection or a regression adjustment using a local linear regression. For each method, values are presented for two different tolerance rates. *n*_1_ to *n*_4_ - relative population sizes of the extant populations, *na*_1_ and *na*_2_ - relative population sizes of the ancestral populations, *rs*_1_ and *rs*_2_ - relative time of the recent split events, *δ_s_* - relative time interval between *rs* and the ancient split event (*t_As_*), *ε_pool_* - experimental error introduced by the pooling procedures, *ε_seq_* - error associated with sequencing and mapping errors, *m_cw_*_1_ and *m_cw_*_2_ - probability per generation that an individual migrates from the Crab population to the Wave population in the first and second locations, respectively (forward in time), *m_wc_*_1_ and *m_wc_*_2_ - probability per generation that an individual migrates from the Wave population to the Crab population in the first and second locations, respectively (forward in time), *m_cc_* - probability per generation that an individual migrates from one Crab population to the other (forward in time), *m_ww_* - probability per generation that an individual migrates from one Wave population to the other (forward in time), *m_aa_* - probability per generation that an individual migrates from one ancestral population to the other (forward in time), *P_cw_* - proportion of the simulated loci where no migration occurs from the Crab to the Wave population; *P_wc_* - proportion of the simulated loci where no migration occurs from the Wave to the Crab population, *P_no_* - proportion of the simulated loci where no migration occurs between ecotypes, 4*N*_2_*m_cw_*_1_ and 4*N*_1_*m_wc_*_1_ - average number of immigrants per generation from Crab to Wave and from Wave to Crab (respectively) in the first location, 4*N*_4_*m_cw_*_2_ and 4*N*_3_*m_wc_*_2_ - average number of immigrants per generation from Crab to Wave and from Wave to Crab (respectively) in the second location, 4*N*_1_*m_cc_*_1_ and 4*N*_3_*m_cc_*_2_ - average number of immigrants per generation from the Crab population in the second location to the first and vice-versa (respectively), 4*N*_2_*m_ww_*_1_ and 4*N*_4_*m_ww_*_2_ - average number of immigrants per generation from the Wave population in the second location to the first and vice-versa (respectively), 4*NA*_1_*m_aa_*_1_ and 4*NA*_2_*m_aa_*_2_ - average number of immigrants per generation from *na*_2_ to *na*_1_ and vice-versa (respectively).

| parameter | REJECTION |  |  |  |  |  | REGRESSION |  |  |  |  |  |
| --- | --- | --- | --- | --- | --- | --- | --- | --- | --- | --- | --- | --- |
|  | tolerance of 0.005 |  |  | tolerance of 0.01 |  |  | tolerance of 0.005 |  |  | tolerance of 0.01 |  |  |
|  | mode | median | mean | mode | median | mean | mode | median | mean | mode | median | mean |
| $n_1$ | 1.26 | 0.66 | 0.58 | 1.39 | 0.70 | 0.62 | 0.18 | 0.17 | 0.18 | 0.18 | 0.17 | 0.18 |
| $n_2$ | 1.25 | 0.67 | 0.60 | 1.35 | 0.69 | 0.61 | 0.20 | 0.19 | 0.20 | 0.19 | 0.17 | 0.18 |
| $n_3$ | 1.20 | 0.64 | 0.58 | 1.29 | 0.66 | 0.60 | 0.19 | 0.18 | 0.19 | 0.19 | 0.18 | 0.19 |
| $n_4$ | 1.26 | 0.68 | 0.60 | 1.40 | 0.71 | 0.63 | 0.20 | 0.19 | 0.20 | 0.21 | 0.20 | 0.21 |
| $na_1$ | 1.99 | 1.13 | 0.98 | 1.97 | 1.16 | 0.99 | 1.91 | 0.99 | 0.89 | 1.92 | 1.02 | 0.90 |
| $na_2$ | 1.92 | 1.10 | 0.98 | 1.97 | 1.13 | 0.99 | 1.86 | 1.00 | 0.91 | 1.91 | 1.03 | 0.92 |
| $rs_1$ | 1.23 | 0.86 | 0.83 | 1.20 | 0.85 | 0.84 | 1.01 | 0.69 | 0.63 | 1.06 | 0.68 | 0.64 |
| $rs_2$ | 1.21 | 0.85 | 0.84 | 1.28 | 0.88 | 0.86 | 1.07 | 0.73 | 0.68 | 1.09 | 0.72 | 0.66 |
| $\delta_s$ | 1.66 | 0.77 | 0.79 | 1.69 | 0.78 | 0.80 | 0.85 | 0.62 | 0.57 | 0.82 | 0.59 | 0.55 |
| $\epsilon_{pool}$ | 0.57 | 0.34 | 0.34 | 0.72 | 0.39 | 0.39 | 0.09 | 0.09 | 0.09 | 0.10 | 0.09 | 0.09 |
| $\epsilon_{seq}$ | 0.22 | 0.18 | 0.23 | 0.27 | 0.24 | 0.29 | 0.02 | 0.02 | 0.02 | 0.02 | 0.02 | 0.02 |
| $m_{cw1}$ | 1.89 | 0.88 | 0.91 | 2.07 | 0.90 | 0.92 | 0.66 | 0.50 | 0.49 | 0.61 | 0.47 | 0.47 |
| $m_{cw2}$ | 2.01 | 0.90 | 0.92 | 2.23 | 0.90 | 0.93 | 0.62 | 0.46 | 0.45 | 0.63 | 0.48 | 0.47 |
| $m_{wc1}$ | 2.04 | 0.88 | 0.91 | 2.02 | 0.90 | 0.92 | 0.64 | 0.48 | 0.47 | 0.60 | 0.45 | 0.45 |
| $m_{wc2}$ | 2.05 | 0.88 | 0.91 | 2.16 | 0.91 | 0.93 | 0.68 | 0.50 | 0.49 | 0.60 | 0.47 | 0.47 |
| $m_{cc}$ | 1.14 | 0.60 | 0.65 | 1.08 | 0.62 | 0.67 | 0.21 | 0.21 | 0.22 | 0.21 | 0.20 | 0.21 |
| $m_{ww}$ | 1.06 | 0.60 | 0.65 | 1.20 | 0.63 | 0.68 | 0.23 | 0.21 | 0.22 | 0.22 | 0.21 | 0.22 |
| $m_{aa}$ | 3.05 | 1.00 | 1.00 | 2.98 | 1.00 | 1.00 | 3.27 | 1.11 | 1.05 | 3.17 | 1.05 | 1.02 |
| $P_{cw}$ | 2.79 | 1.00 | 0.94 | 3.04 | 0.99 | 0.96 | 2.16 | 0.90 | 0.79 | 2.30 | 0.91 | 0.81 |
| $P_{wc}$ | 2.82 | 0.98 | 0.94 | 3.02 | 1.01 | 0.97 | 2.15 | 0.89 | 0.78 | 2.36 | 0.89 | 0.80 |
| $P_{no}$ | 0.34 | 0.20 | 0.16 | 0.33 | 0.20 | 0.16 | 0.25 | 0.13 | 0.11 | 0.22 | 0.11 | 0.10 |
| $4N_2m_{cw1}$ | 1.35 | 0.87 | 0.75 | 1.38 | 0.87 | 0.76 | 0.37 | 0.33 | 0.32 | 0.34 | 0.31 | 0.31 |
| $4N_1m_{wc1}$ | 1.36 | 0.87 | 0.75 | 1.39 | 0.90 | 0.77 | 0.36 | 0.33 | 0.33 | 0.35 | 0.31 | 0.31 |
| $4N_4m_{cw2}$ | 1.34 | 0.86 | 0.74 | 1.43 | 0.90 | 0.77 | 0.36 | 0.33 | 0.33 | 0.37 | 0.33 | 0.33 |
| $4N_3m_{wc2}$ | 1.31 | 0.85 | 0.74 | 1.36 | 0.88 | 0.76 | 0.34 | 0.31 | 0.31 | 0.34 | 0.31 | 0.31 |
| $4N_1m_{cc1}$ | 1.08 | 0.78 | 0.70 | 1.15 | 0.81 | 0.70 | 0.30 | 0.26 | 0.25 | 0.34 | 0.27 | 0.25 |
| $4N_3m_{cc2}$ | 1.09 | 0.78 | 0.67 | 1.15 | 0.80 | 0.69 | 0.30 | 0.26 | 0.25 | 0.34 | 0.28 | 0.27 |
| $4N_2m_{ww1}$ | 1.06 | 0.77 | 0.67 | 1.08 | 0.78 | 0.69 | 0.32 | 0.27 | 0.26 | 0.32 | 0.27 | 0.26 |
| $4N_4m_{ww2}$ | 1.09 | 0.78 | 0.68 | 1.14 | 0.81 | 0.70 | 0.35 | 0.29 | 0.28 | 0.32 | 0.27 | 0.27 |
| $4NA_1m_{aa1}$ | 1.72 | 1.17 | 0.99 | 1.69 | 1.17 | 1.00 | 1.72 | 1.10 | 0.96 | 1.69 | 1.09 | 0.94 |
| $4NA_2m_{aa2}$ | 1.70 | 1.15 | 0.99 | 1.72 | 1.16 | 0.99 | 1.69 | 1.07 | 0.95 | 1.71 | 1.09 | 0.96 |

**Table S4:** Estimates for relative parameters of *Littorina saxatilis* populations. Parameter inference was performed with a regression adjustment using a local linear regression and a tolerance rate of 0.01. Results are shown for the Arsklovet and Burela populations using the parallel origin scenario. For this model, *n*_1_ and *n*_2_ correspond to the Arsklovet Crab and Wave population respectively, while *n*_3_ and *n*_4_ correspond to the Burela Crab and Wave population respectively. For each parameter, the value outside brackets corresponds to the mean of the posterior distribution and in-between brackets is the 95% credible interval. Parameters indicated here are the same as in table S2.

| parameter | relative estimate |
| --- | --- |
| $n_1$ | 0.200 (0.126 - 0.322) |
| $n_2$ | 0.259 (0.145 - 0.455) |
| $n_3$ | 0.830 (0.344 - 1.727) |
| $n_4$ | 0.574 (0.239 - 1.157) |
| $na_1$ | 1.079 (0.139 - 2.709) |
| $na_2$ | 1.307 (0.158 - 2.799) |
| $rs_1$ | 0.288 (0.026 - 1.130) |
| $rs_2$ | 0.797 (0.105 - 2.363) |
| $\delta_s$ | 1.934 (0.309 - 2.925) |
| $m_{cw1}$ | 0.00057 (0.00014 - 0.00096) |
| $m_{wc1}$ | 0.00065 (0.00019 - 0.00097) |
| $m_{cw2}$ | 0.00020 (0.00002 - 0.00070) |
| $m_{wc2}$ | 0.00042 (0.00007 - 0.00089) |
| $m_{cc}$ | 0.00004 (0.00002 - 0.00007) |
| $m_{ww}$ | 0.00001 (0.00000 - 0.00003) |
| $m_{aa}$ | 0.00003 (0.00000 - 0.00009) |
| $P_{cw}$ | 0.024 (0.000 - 0.172) |
| $P_{wc}$ | 0.028 (0.000 - 0.186) |
| $P_{no}$ | 0.037 (0.005 - 0.088) |

**Figure S1:**
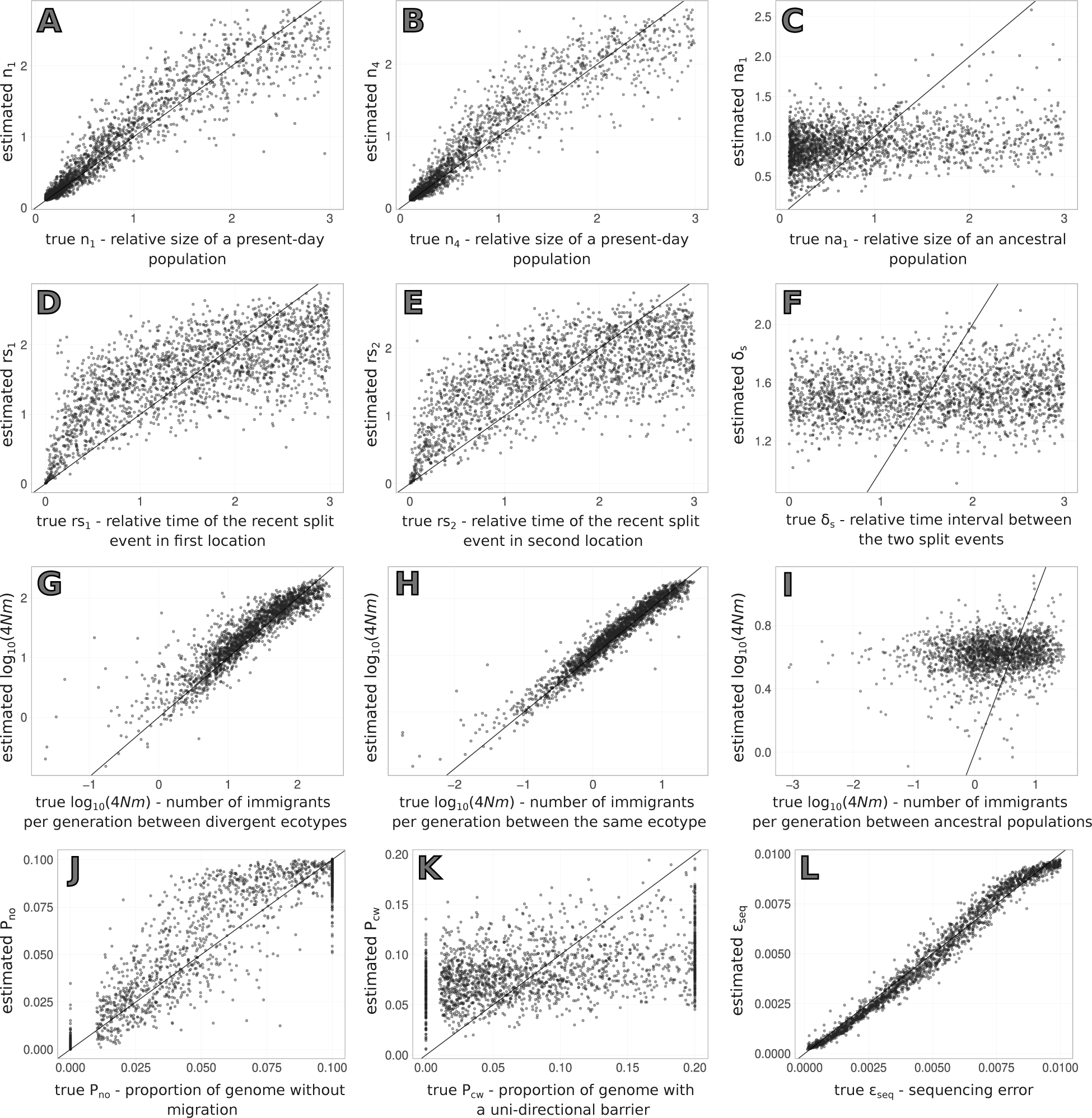
Results of the cross-validation for parameter estimation using the parallel origin scenario. The y-axis displays the estimated values, plotted against the true parameter values on the x-axis. Estimates correspond to the mean of the posterior obtained with a tolerance rate of 0.01. Parameters shown here are: A and B - relative size of a present-day population (*n*_1_ and *n*_2_); C - relative size of an ancestral population (*na*_1_); D and E - relative time of the recent split event in the first (*rs*_1_) and second locations (*rs*_2_) respectively; F - relative time interval between the two split events (*δ_s_*); G - average number of immigrants per generation in *log*_10_ scale between populations of the divergent ecotypes; H - average number of immigrants per generation in *log*_10_ scale between populations of the same ecotype; I - average number of immigrants per generation in *log*_10_ scale between ancestral populations; J - proportion of the simulated loci where no migration occurs between ecotypes (*P_no_*); K - proportion of the simulated loci where no migration occurs from one of the ecotypes to the other (*P_cw_*) and L - sequencing error.

**Figure S2:**
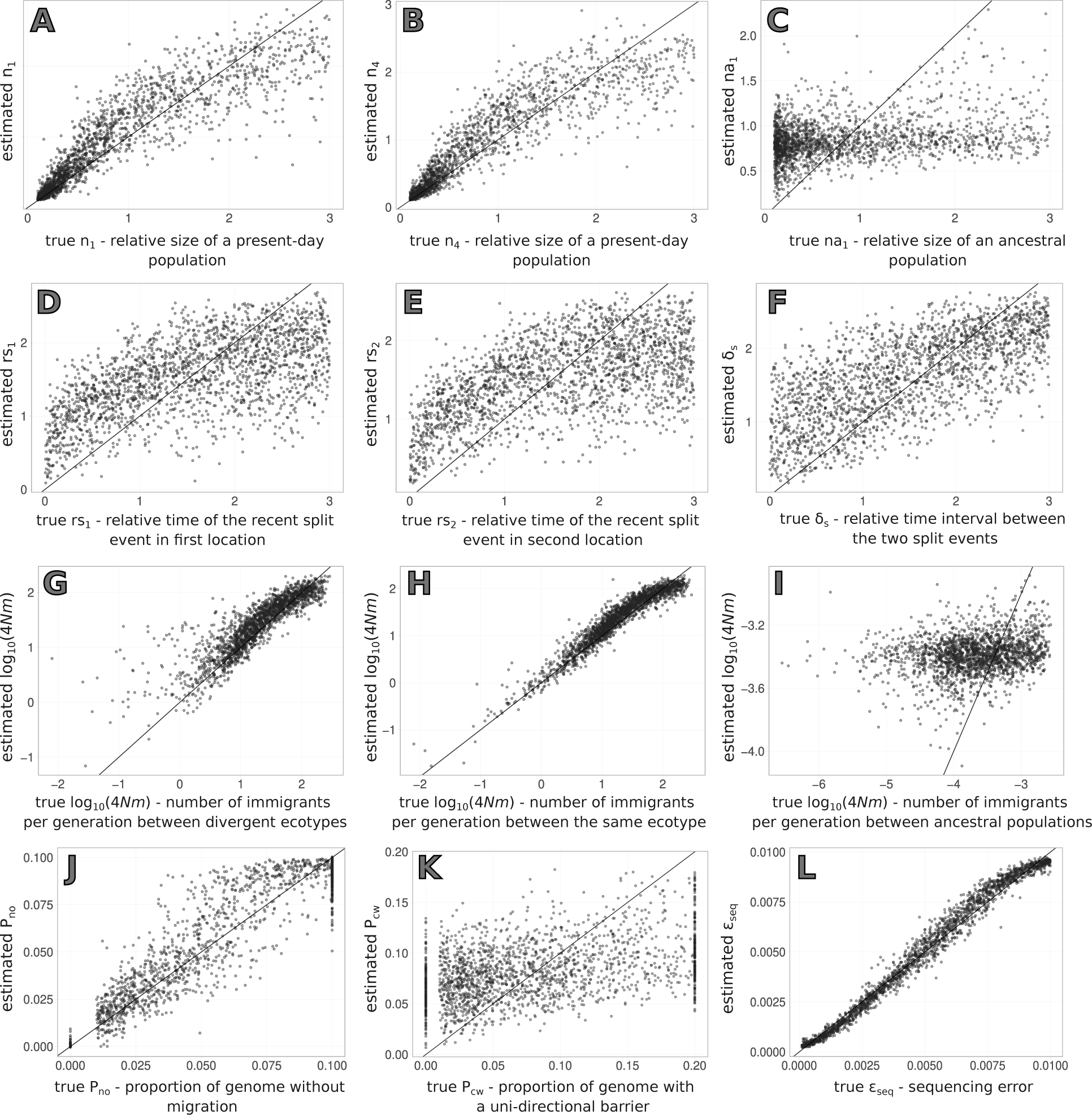
Results of the cross-validation for parameter estimation using the single origin scenario. The y-axis displays the estimated values, plotted against the true parameter values on the x-axis. Estimates correspond to the mean of the posterior obtained with a tolerance rate of 0.01. Parameters shown here are: A and B - relative size of a present-day population (*n*_1_ and *n*_2_); C - relative size of an ancestral population (*na*_1_); D and E - relative time of the recent split event in the first (*rs*_1_) and second locations (*rs*_2_) respectively; F - relative time interval between the two split events (*δ_s_*); G - average number of immigrants per generation in *log*_10_ scale between populations of the divergent ecotypes; H - average number of immigrants per generation in *log*_10_ scale between populations of the same ecotype; I - average number of immigrants per generation in *log*_10_ scale between ancestral populations; J - proportion of the simulated loci where no migration occurs between ecotypes (*P_no_*); K - proportion of the simulated loci where no migration occurs from one of the ecotypes to the other (*P_cw_*) and L - sequencing error.

**Figure S3:**
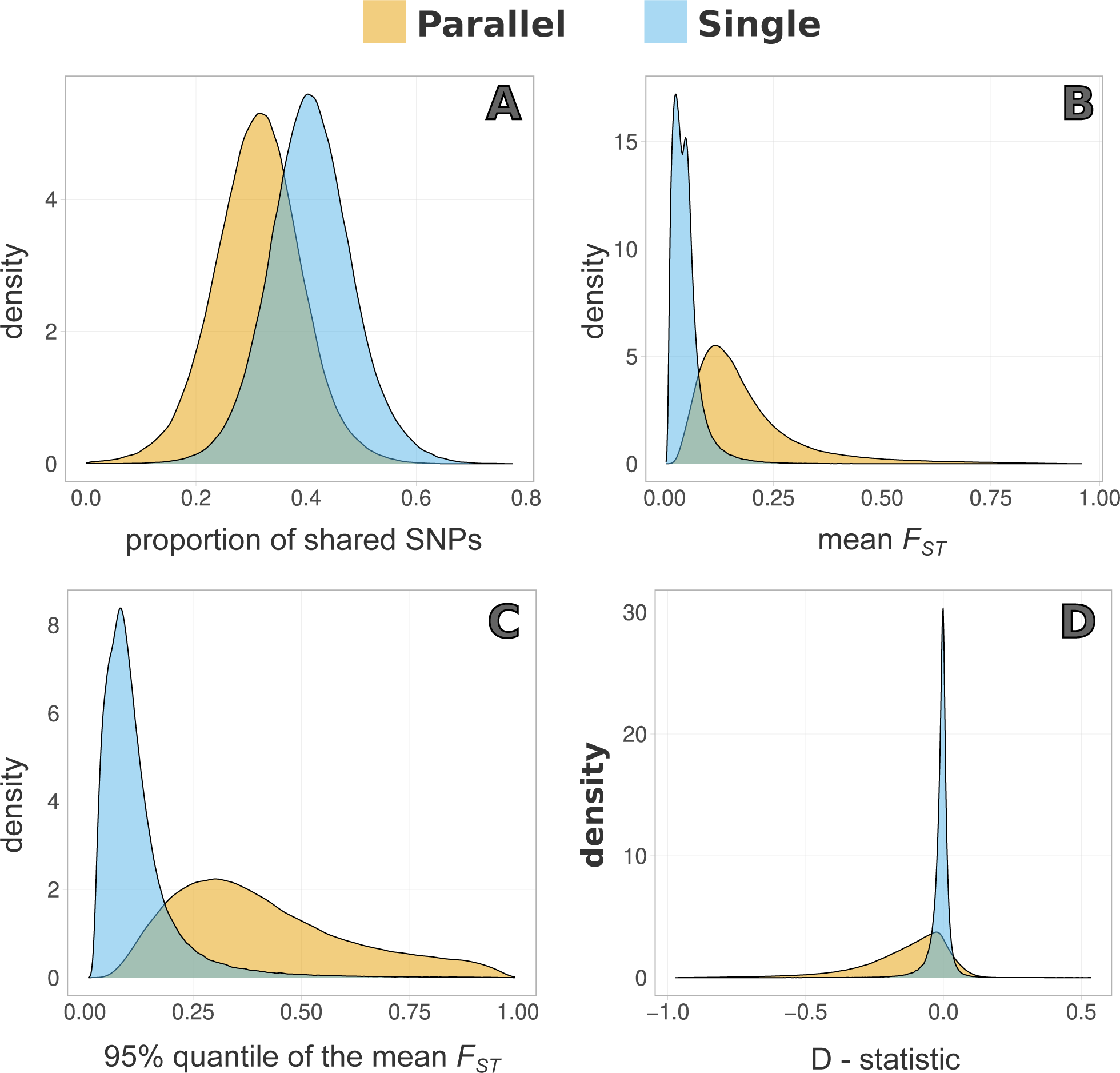
Distribution of summary statistics obtained for the single and parallel origin scenarios. Summary statistics are: A - proportion of shared SNPs between all four present-day populations, B - mean pairwise *F_ST_* between Wave populations at different locations, C - 95% quantile of the mean pairwise *F_ST_* between Crab populations at different locations and D - D-statistic.

